# Ferroptosis of selective progenitor cell types during embryogenesis ensures proper organ morphogenesis

**DOI:** 10.64898/2026.11.03.764065

**Authors:** Montasir Elahi, Shicong Song, Wei-Bin Shen, Anna Wu Chen, Md Mahadi Hassan, Mohammad Sazzadul Islam, E. Albert Reece, Robin Roychaudhuri, Guanglei Wang, Lihua Zeng, Bingbing Wang, Jin-Gu Lee, Ivy Han, Qitao Ran, Liang Li, Shuan Zhao, Irina Burd, Dana Marie Roque, Hangnoh Lee, Peixin Yang

**Author notes:** Correspondence: Peixin Yang, PhD.; and Wie-Bin Shen.edu; and Montasir Elahi, Department of Obstetrics, Gynecology & Reproductive Sciences, University of Maryland School of Medicine, 655 W. Baltimore Street, Baltimore, MD, 21201. These authors contribute equally to this work (Montasir Elahi, Shicong Song, Wei-Bin Shen).

## Abstract

Ferroptosis, a form of non-apoptotic cell death, is induced by lipid peroxidation resulting from an intracellular accumulation of labile iron^1^. While apoptosis is involved in embryonic development^2^, the role of ferroptosis in embryogenesis is unknown. Here, we report that the ferroptosis process occurs in the embryonic lung and kidney at mid- and late-gestation. In these tissues, pro-ferroptotic cells include subpopulations of the Sox9^+^ distal lung epithelial cells and the Flk1^+^ vascular progenitors in the kidney glomeruli. Lipidomic analysis reveals mixed lipid peroxidation as the driver for ferroptosis. The role of ferroptosis in the morphogenesis of these organs was confirmed by triple positivity of TfR1, 4-HNE and TUNEL, mitochondrial defects, cell membrane breakage, dead cell remnants, single-cell RNA-sequencing of a ferroptosis gene signature. Genetic manipulation of *Gpx4*, an endogenous inhibitor of ferroptosis. Further, genetic knockout or transgenic overexpression of *Gpx4* affects lung and kidney development and results in cystic lungs and kidneys. Notably, conditional deletion *Gpx4* results in hypoplasia of the lung epithelia and of the renal glomeruli, while transgenic overexpression of *Gpx4* resulted in atelectasis with collapsed lung alveoli and hypertrophic glomeruli. Together, these findings show that ferroptosis regulates lung and kidney morphogenesis by acting in key progenitor cell populations during gestation. Supplementation of iron or polyunsaturated fatty acid during pregnancy has implications for organogenesis.

## Introduction

Ferroptosis, a form of programmed cell death, is induced when intracellular labile iron accumulates, leading to lipid peroxidation^1^. Cells take up ferric (Fe^3+^)-bound transferrin (Tf) via the cell membrane Tf receptor (TfR1)^3^, which is recently recognized as a possible tissue marker for ferroptosis^4^. Increasing cellular iron import through the Tf-TfR1 system can trigger ferroptosis^5^. In contrast, reducing intracellular iron levels by enhancing its export results in ferroptosis resistance^6,7^. It has been shown that maternal iron deficiency in early embryonic development leads to cardiovascular developmental defects in the offspring^8^, suggesting that iron-mediated ferroptosis is required for the intricate processes of embryonic organ morphogenesis. Cellular labile iron initiates and propagates lipid peroxidation through the non-enzymatic Fenton reaction^1^. Iron-dependent lipid peroxidation is an essential element in various pathways or mechanism-induced ferroptosis^9^. Although ferroptosis implies that iron is the key, the role of iron in ferroptosis is far from being understood. It remains to be firmly established that Fenton chemistry is actually present in cells and tissues causing ferroptosis.

The hallmark of ferroptosis, lipid peroxidation, mainly occurs in the polyunsaturated fatty acyl (PUFA) moieties of phospholipids in the lipid bilayer of the cell membrane^10^. During ferroptosis, reactive oxygen species (ROS) generated by cellular metabolic reactions can directly oxidize PUFAs, leading to the formation of advanced lipid peroxidation products, including 4-hydroxynonenal (4-HNE) and malondialdehyde (MDA)^1^. The expression of the lipid metabolic enzyme acyl-CoA synthetase long-chain family member 4 (Acsl4) also positively regulates ferroptosis by increasing PUFA incorporation into membrane phospholipids^11^. Thus, the amount of PUFAs in membrane lipids is proportionately linked to ferroptosis sensitivity in cells^1^. While lipid peroxidation-induced ferroptosis has been involved in pathogenic conditions^12^, such a role has not been demonstrated in cells of the developing embryo. Intrauterine transfer of PUFAs is important for proper fetal growth and development^13^. Additionally, the *acsl4* gene is essential for zebrafish embryonic development^14^. Such evidence suggests the potential role of lipid peroxidation-induced ferroptosis in mammalian embryogenesis.

The enzyme glutathione peroxidase 4 (GPX4) inhibits ferroptosis by reducing oxidized phospholipids to corresponding phospholipid alcohols^10^. The selective ferroptosis inducer RAS-selective lethal 3 (RSL3) induces ferroptosis through GPX4 inhibition^15^. Other antioxidant pathways, including radical trapping antioxidants and the ferroptosis suppressor protein 1 (FSP1)-initiated pathway, block ferroptosis by reducing lipid peroxidation^1,16^. Conditional knockout of the *Gpx4* gene causes lipid peroxidation and consequent ferroptosis in mice^17,18^, while *Gpx4* germline deletion confers lethality in midgestation^19^. The spatial and temporal changes of GPX4 expression during embryonic development^20^ suggest an important role of this lipid peroxidation detoxifying enzyme in embryogenesis.

In the present study, we used the two ferroptosis markers TfR1 and 4-HNE, along with the TUNEL (terminal deoxynucleotidyl transferase dUTP nick end labeling) labeling of dead cells, to longitudinally and systemically define ferroptotic cells in the developing embryo in mice. We found that a subpopulation of Sox9^+^ lung distal epithelium and a group of Flk1^+^ (Fetal liver kinase 1) vascular progenitors in the renal glomeruli as ferroptotic cells. We also found that a mixed type of lipid peroxidation is an inducer of ferroptosis in the developing lung and the kidney. Loss- and gain-of-function of the ferroptosis inhibitor GPX4 in the lung or the kidney altered the morphogenesis of these two organs. Notably, loss of *Gpx4* in the lung or kidney altered the morphogenesis of these two organs, leading to cystic structure formation, while overexpression of *Gpx4* in the lung or the kidney led to atelectasis with collapsed lung alveoli and hypertrophic glomerulus with the loss of the Bowman’s space, respectively. These findings underscore that ferroptosis is required for proper embryonic morphogenesis of the lung and the kidney.

## Results

### Identification of ferroptosis markers-positive cells in specific organs of the embryo

To reveal potential ferroptotic cells during embryogenesis, we surveyed the developing embryo for ferroptotic cells using a TfR1 antibody from embryonic day 8.5 (E8.5) to E18.5, which covers the early middle-to-late periods for organogenesis and morphogenesis (**Fig. 1a-c**). TfR1-positive (TfR1^+^) cells were detected in the lung from E10.5 until E16.5 (**Fig. 1a, b, Fig. S1a**), in the eye at E13.5 but not at E16.5 (**Fig. 1a**), in the pancreas at E13.5 and E16.5 (**Fig. 1a**) and in the kidney from E16.5 to P1 (**Fig. 1a, c**). In the *TfR1* conditional knockout (cKO) E14.5 lungs or E14.5 lungs treated with Ferrostatin-1, an ferroptosis inhibitor, TfR1^+^ cells were diminished whereas the ferroptosis activator Erastin triggered TfR1-positive staining in E12.5 lung (**Fig. S1b**). In the lung, TfR1^+^ cells were localized to tubular structures (**Fig. 1b**). In the kidney, TfR1^+^ cells resided in capsule-like structures, particularly in well-formed glomeruli (**Fig. 1c**). In the eye, TfR1^+^ cells were present in the most inner epithelial cells of the eyeball (**Fig. 1a**), and in the pancreas TfR1^+^ cells were scattered throughout the entire organ (**Fig. 1a**).

**Figure 1.**
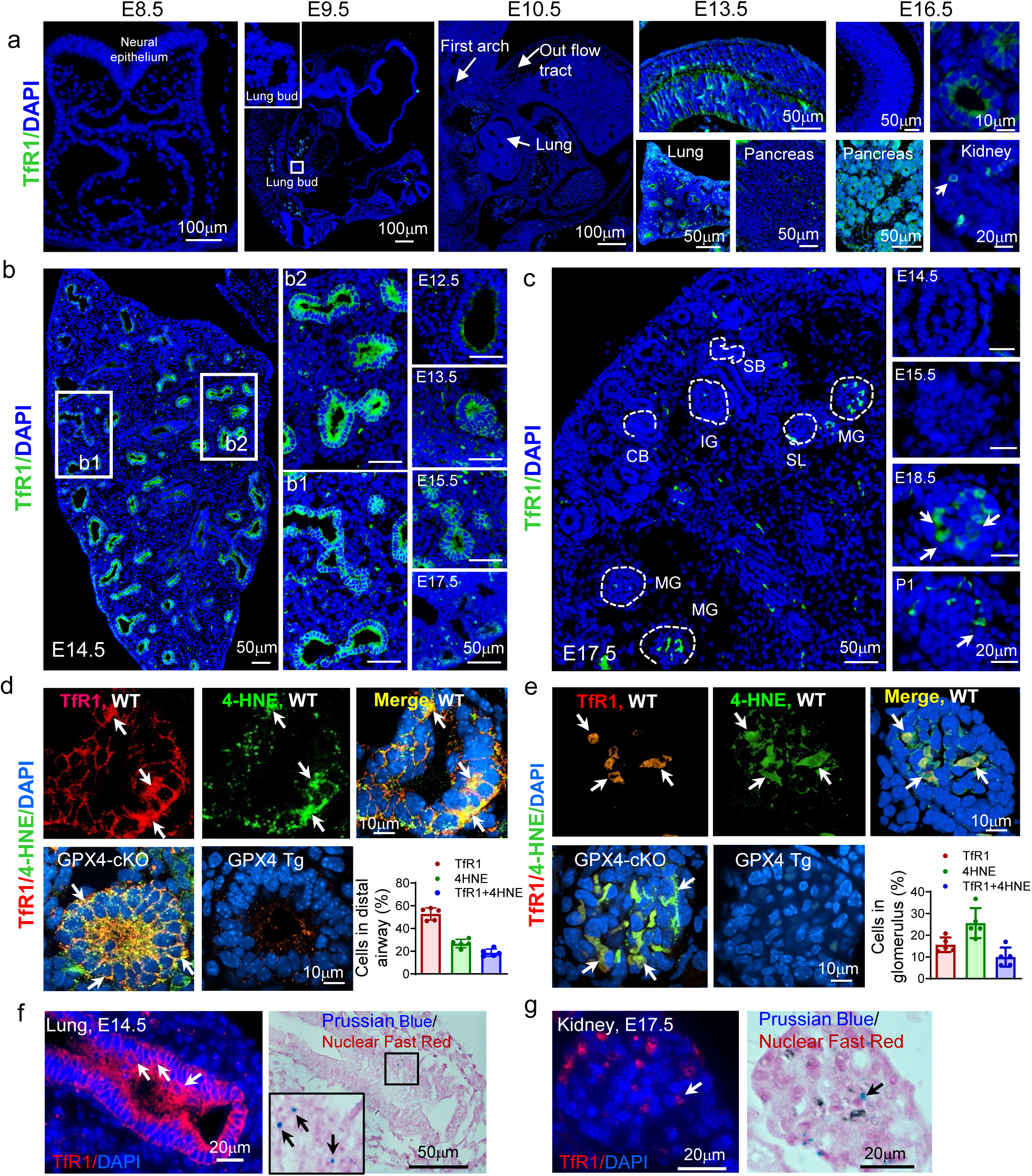
The pro-ferroptosis process is present in multiple organs during development. Representative images of TfR1 staining in different organs at different embryonic stages (**a**). Representative images of TfR1 staining in lung (**b**) and kidney (**c**) at different embryonic stages. Identification of TfR1- and 4-HNE-double-positive cells in E14.5 lungs when the TfR1 expression peaks (**d**) and E18.5 kidney (**e**) in wild-type, GPX4 Tg, and GPX4 cKO embryos; white arrows indicate the double positive cells where both 4-HNE and TfR1 showed a pattern of membrane staining. Quantification of the percentage of single TfR1, 4-HNE and TfR1 and 4-HNE double-positive cells in the distal airways of E14.5 lung and glomeruli of E18.5 kidney in the bar graphs (lower right panels of **d** and **e**). Embryos from five litters (*n* = 5) each group were analyzed. One embryo from each litter and 3 sections per embryo were used in immunofluorescence staining. For statistical analysis, one-way ANOVA followed by *Tukey* test was used. A positive control of the ferroptosis GPX4 cKO (GPX4f/f;Nkx2.1-Cre) for lungs and GPX4 cKO (GPX4f/f:Flk1-Cre) for kidneys and a negative control GPX4 Tg E14.5 lungs and E18.5 lungs in the lower panels **(d**, **e)**. Representative images TfR1 protein staining and Prussian blue iron staining in E14.5 lung distal epithelium (**f**) and E18.5 kidney glomeruli (**g**) in serial adjunct sections; white arrows indicate the presence of the iron in TfR1 positive cells. Embryos from three litters (*n* = 3) each group were analyzed. Two to three embryos from each litter and 3 sections per embryo were stained. SB, S-shaped body; CB, Common shaped body; IG, Intermediate mature glomerulus; SL, Single capillary loop; MG, Mature glomerulus. Scale bars are presented.

Next, we co-stained sections for TfR1 and the lipid peroxidation marker 4-HNE (4-Hydroxynonenal) (**Fig. 1d, Fig. S1c**) in the lung and found that 18.6% of the TfR1^+^ cells are 4-HNE positive (**Fig. 1d**). 4-HNE positive staining showed dot-like cell membrane signals and was abrogated in E14.5 lung epithelia when the antibody was neutralized by BSA-4-HNE prior to staining (**Fig. S1c**). Additionally, E14.5 *Tfr1* null lungs or Ferrostatin-1-treated wild-type (WT) lungs did not show any 4-HNE positive staining whereas Erastin enhanced 4-HNE staining signal (**Fig. S1d**), confirming the specificity of the 4-HNE antibody. In the developing kidney, 9.7% of TfR1^+^ cells were also 4-HNE positive in the glomeruli (**Fig. 1e, Fig. S1e**). However, E18.5 TfR1 null kidneys or WT lungs treated with Ferrostatin-1 did not have any double TfR1 and 4-HNE staining positivity in E18.5 kidneys whereas Erastin induced excessive double positive cells in E15.5 kidneys (**Fig. S1f**). This evidence indicates that these double-positive cells possess two key characteristics of ferroptosis: increased iron cellular uptake via increased TfR1 and the co-presence of lipid peroxidation. Indeed, ferric ion (Fe^3+^)-enriched cells with TfR1 positivity were identified in both lung and kidney (**Fig. 1f, g**).

### Ferroptosis markers^+^ cells exhibit the features of lytic cell death

In E14.5 lungs, a few of the TfR1 and 4-HNE double-positive cells were TUNEL positive (**Fig. 2a**), and these triple-positive cells were diminished in GPX4 Tg lungs but increased in GPX4 cKO lungs (**Fig. 2a**). A comparable phenomenon was also observed regarding TUNEL positivity of TfR1 and 4-HNE double-positive cells in the developing glomeruli (**Fig. 2b**). The glomerular cells from GPX4 Tg mice did not exhibit TUNEL positivity but GPX4 cKO enhanced TUNEL positivity in E18.5 glomeruli (**Fig. 2b**). The ferroptosis inducer Erastin triggered the emergence of triple-positive TfR1, 4-HNE and TUNEL cells in E12.5 lungs and E15.5 kidney (**Fig. S2a**), embryonic stages when these two organs did not otherwise exhibit positivity of TfR1 and 4-HNE (**Fig. 1**). Ferrostatin-1 abolished the presence of these triple-positive cells in E14.5 lungs and E18.5 kidney (**Fig. S2a**), stages when the double-positivity of TfR1 and 4-HNE reached a plateau under WT conditions (**Fig. 1**).

**Figure 2.**
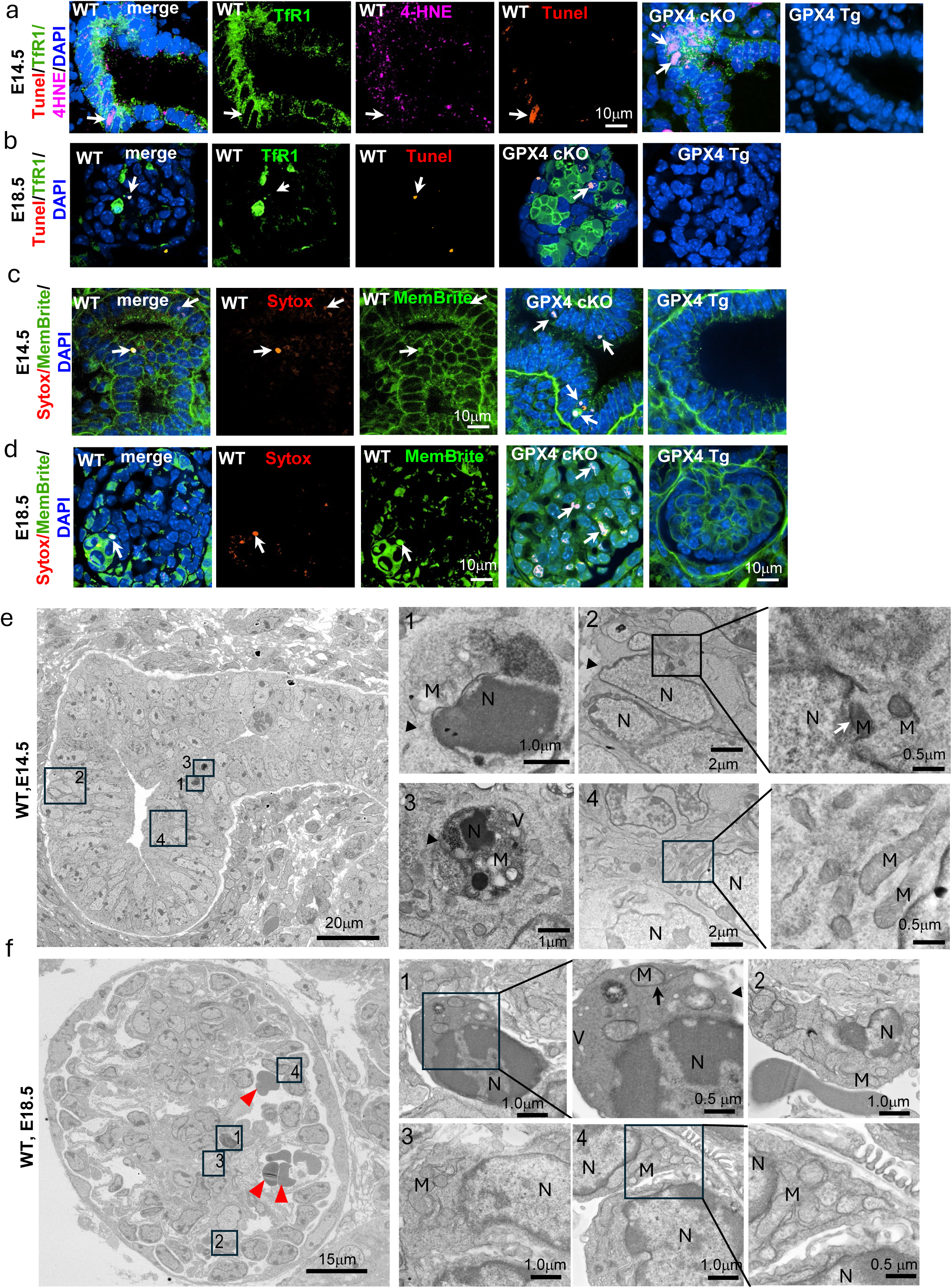
Identification of cell death in TfR1 and 4-HNE double positive cells in developing lung distal epithelium and kidney glomeruli. TUNEL positivity in TfR1 and 4-HNE double positive cells in E14.5 lungs (**a**) and E18.5 kidney (**b**). Triple TfR1, 4-HNE and TUNEL positive cells in wild-type (WT), GPX4 cKO (GPX4f/f;Nkx2.1-Cre) for lungs and GPX4 cKO (GPX4f/f:Flk1-Cre) for kidneys, and GPX4 Tg. Triple positive cells are indicated by white arrows. Representative images of E14.5 lung **(c)** and E18.5 kidney **(d)** in cell membrane dye (MemBrite) and nuclear dye (Sytox) perfusion staining. Double positive cells are indicated with white arrows. Lung or kidney of 3 embryos from three different litters (*n* = 3) were analyzed, minimum 3 sections per embryo were stained. Electron microscopic images for E14.5 lung distal epithelium **(e)** and E18.5 kidney glomerulus **(f).** The presence of a dead cell reminiscence (e-1), condensed nucleus (N), swollen lytic mitochondria (M) and the damaged membrane indicated by black arrowheads (e-1, e-2). A cell with condensed and shrinked mitochondria with abnormal cristae, compromised membrane indicated by white arrow (e-2). Shrinked lytic mitochondria, in a dead cell (e-3), condensed nucleus (N), lytic mitochondria (M) and stressed vacuole (V) and the damaged membrane indicated by black arrowhead. Mitochondria in the apparently healthy cell in the lung (e-4). In E18.5 glomerulus, a dying cell (f-1), cells with mitochondrial defects (f-2, f-3) and an apparently healthy cell (f-4) are indicated. Presence of the blood cells are indicated by red arrowheads, and M: mitochondria and N: Nucleus. In the dying cell (f-1) condensed nucleus (N), swollen mitochondria (M) and stressed vacuole (V) and the compromised membrane are indicated by black arrowhead. Mitochondria defective cells showed both shrinked (f-2) and swollen mitochondrial morphology (f-3). Embryos from three different litters (*n* = 3) were analyzed.

We perfused lung and kidney tissues with a mixture of Sytox Red and Membrite membrane dyes. In the lung, a few of Sytox Red-labeled nuclei were observed in the distal epithelium where triple-positive TfR1, 4-HNE and TUNEL cells resided (**Fig. 2c**), indicating these cells have damaged cell membranes. We saw nucleus dye positivity in E18.5 glomeruli (**Fig. 2d**). In cells from GPX4 cKO mice we found enhanced cell nucleus dye penetration whereas in cells from GPX4 Tg mice such penetration was abrogated (**Fig. 2c, d**). We further found that Erastin promoted nucleus dye penetration in cells of E12.5 lungs and E15.5 kidney whereas Ferrostatin-1 abolished nucleus dye penetration in cells of E14.5 lungs and E18.5 kidney (**Fig. S2b**). By election microscopy (EM) we found evidence of dying cells and cells with mitochondrial defects and cell membrane damage in the lung distal epithelium and in glomeruli at E14.5 and E18.5, respectively (**Fig. 2e, f**), while the numbers of these dying cells and cells with mitochondrial alterations and cell membrane defects were lower in lung and kidney tissues from GPX4 Tg mice (**Fig. S2c**) and higher in the glomeruli of GPX4 cKO mice (**Fig. S2d**). Utilizing TfR1 antibody-mediated pulldown assays in lung and kidney cells, we found mitochondrial defects but such defects were absent in flowthrough cells that were TfR1-negative (**Fig. S2e, f**). These results support the notion that double-positive TfR1 and 4-HNE cells exhibit key features of lytic cell death, including TUNEL positivity and dead cell EM characteristics, as well as mitochondrial and cell membrane damage.

### A population of lung Sox9^+^ distal epithelial cells are pro-ferroptotic

To further characterize the TfR1^+^ cells in the lung, we utilized lung epithelial cell markers in immunofluorescence staining assays. We found that the nuclear staining of Sox9, a lung distal epithelial cell marker^21^, was perfectly complementary with the cell membrane staining of TfR1 (**Fig. 3a, b, c**). On the other hand, epithelial cells in the trachea labeled with Sox2, a recognized marker of trachea epithelium^22^, were TfR1-negative at E14.5 (**Fig. 3a, b, c**). In contrast, Sox9^+^ cells in E10.5 lung buds at the beginning of branching morphogenesis were TfR1negative (**Fig. 3c**). Sox9^+^ cells were also positive for Ecad expression at that developmental stage (**Fig. 3d**). These findings suggest that TfR1^+^ cells in the lung are distal airway epithelial cells (**Fig. 3a, b, c**) and that the number of TfR1^+^ cells reached their peak at E14.5 (**Fig. 1b**).

**Figure 3.**
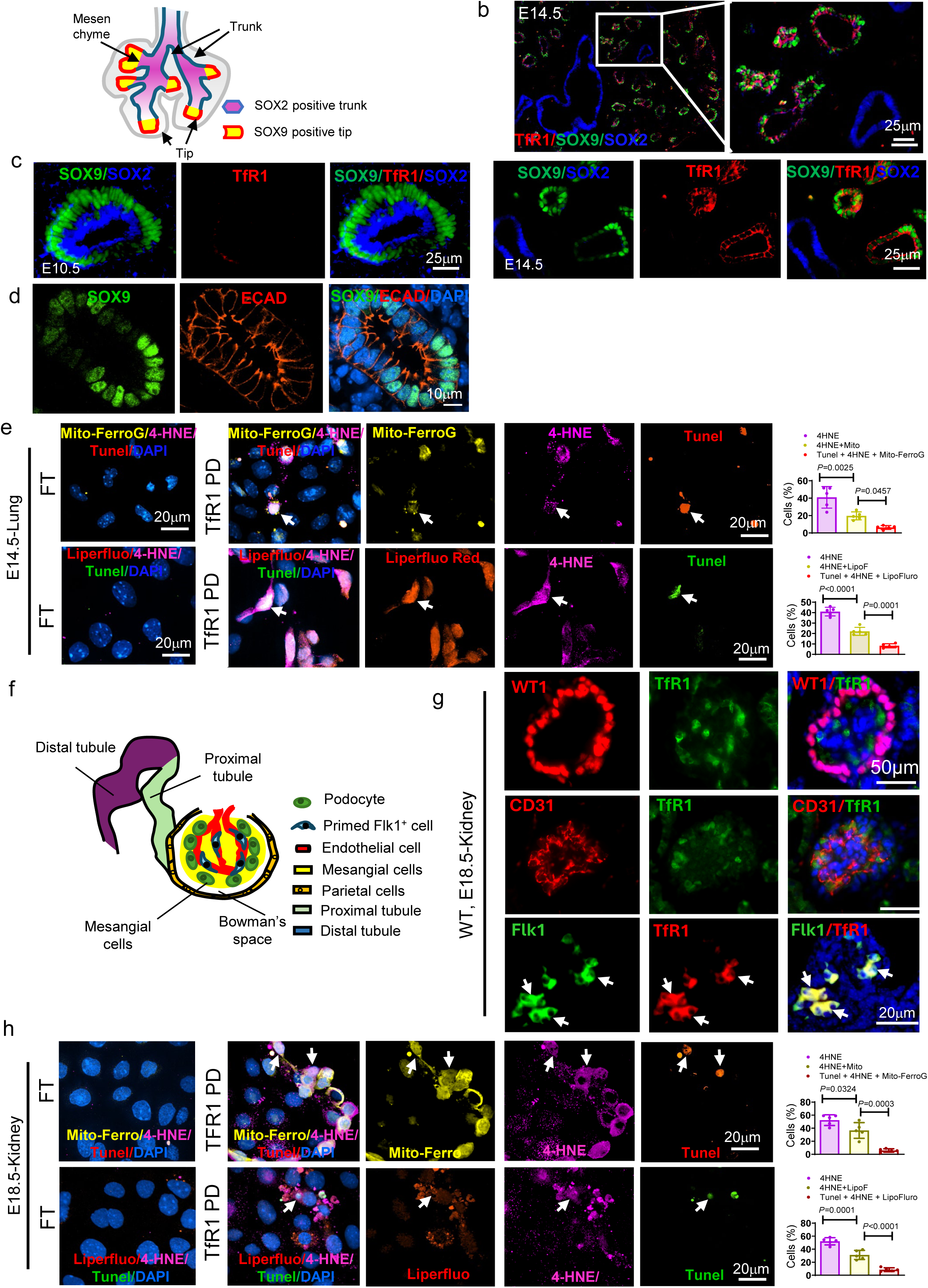
Sox9^+^ epithelial cells in the developing lung and Flk1^+^ cells in the renal glomeruli undergo ferroptosis. (**a**) A schematic illustrating the structure of the E14.5 mouse lung; Sox2^+^ cells are located in the trunk and Sox9^+^ cells are located in the distal epithelium. TfR1^+^ cells are depicted as red, whereas Sox2^+^ cells are shown in blue. Sox2, Sox9 and TfR1 triple staining in E14.5 lung (**b**). Representative images of Sox9, Sox2 and TfR1 staining in E10.5 and E14.5 lungs (**c**). Sox9 and E-cadherin (Ecad) were perfectly overlapping with each other by double staining using E14.5 lung (**d**). TfR1^+^ pulldown and flowthrough cells from E14.5 lung subjected to Mito-FerroGreen (upper) or Liperfluoro (lower) staining with 4-HNE and TUNEL labeling (**e**). Statistic comparisons of double Mito-FerroGreen or Liperfluoro, and 4-HNE positive cells, and double-positive cells with TUNEL positivity are presented in the bar graphs. Lung from the embryos of five different litters (*n* = 5) were subjected for analysis and each litter consists of 8-9 embryo, and the cell samples have triplicate for each experiment. Data were analyzed by One-way ANOVA followed by *Tukey’s* multiple comparison test. (**f**) Graphic structure of E18.5 kidney. (**g**) Representative images of TfR1 staining with podocyte marker WT1, endothelial cell marker CD31, and endothelial progenitor marker Flk1 in E18.5 kidney. Two to three embryos from each litter and four to five sections per embryo were stained. (**h**) Mito-FerroGreen or Liperfluo, 4-HNE and TUNEL triple positive cells in E18.5 kidney TfR1 pulldown and flow-through cells. Percentages of double Mito-FerroGreen or Liperfluo, and 4-HNE positive cells, and double positive cells with TUNEL positivity were presented in the bar graphs (**h**). Kidney from the embryos of five different litters (*n* = 5) were subjected for analysis and each litter consists of 8-9 embryo, and the cell samples have triplicate for each experiment. *P* values from One-way ANOVA followed by *Tukey* test.

To further define whether some or all the TfR1^+^ cells in the lung are pro-ferroptotic cells, we established a single-cell ferroptosis assay. We isolated TfR1^+^ cells by antibody-conjugated magnet beads. We found that ∼40% of the pull-downed cells were positive for 4-HNE (**Fig. 3e**). In these pull-downed TfR1 and 4-HNE double-positive cells, almost half of them (or ∼20% of the total cells) were Mito-FerroGreen-positive (**Fig. 3e**). Mito-FerroGreen, a fluorescent probe which detects ferrous ion (Fe^2+^) in mitochondria, is a ferroptosis detection method for intact living cells^23^. 6.5% of double-positive 4-HNE and Mito-FerroGreen cells in the TfR1 pulldown were TUNEL positive (**Fig. 3e**), suggesting these cells are ferroptotic cells. Additionally, ∼23% of pull-downed TfR1^+^ cells had a higher degree of lipid peroxidation, as assessed by Liperfluo^24^ and these cells were 4-HNE-positive (**Fig. 3e**). Eight percent of double-positive 4-HNE and Liperfluo cells in the TfR1 pulldown were TUNEL positive (**Fig. 3e**). These results indicate that developmental pro-ferroptotic cells exhibit Mito-FerroGreen and Liperfluo positivity. We also treated E14.5 lung tissues *in vitro* with the apoptosis-inducer, camptothecin. Doing so resulted in a marked number of apoptotic cells in lungs as determined by immunofluorescence staining of cleaved caspase-3 antibody (**Fig. S3a**). No apoptotic cells and necroptotic cells were detected in epithelial cells of the embryonic lung with the cleaved caspase 3 antibody and antibodies to the necroptosis markers p-MLKL, p-RIPK1 and p-RIP3, respectively (**Fig. S3a, b**). Rather, we only found evidence for necroptosis in cells from tissues treated with an inducer of this process (**Fig. S3b**). We confirmed the specificity of the necroptosis-related antibodies by neutralization of the antibodies with their corresponding proteins prior to immunostaining (**Fig. S3b**). As there are two pathways that inhibit ferroptosis, we performed FSP1 antibody immunofluorescence staining to determine if FSP1 participates in the regulation of ferroptosis in lung development. We found no FSP1-positive cells in lung distal epithelial (**Fig. S3c**). These findings suggest that ferroptosis of distal epithelial cells, rather than apoptosis or necroptosis, plays a role in lung branching morphogenesis.

### Glomerular Flk1^+^ progenitors are pro-ferroptotic

As noted above, we found that TfR1^+^ cells were first detected in the E16.5 kidney and the number of these cells reached their peak in the kidney at E18.5 (**Fig. 1c, S1e**). TfR1^+^ cells were generally present inside the glomeruli (**Fig. 1e)**. The podocyte marker WT1 (Wilms’ tumor 1)^25^ did not overlap with the TfR1 staining (**Fig. 3f, g**). The TfR1^+^ cells were not endothelial cells because they were not CD31 (cluster of differentiation 31) positive (**Fig. 3g**). However, TfR1^+^ cells were positive for the vascular endothelial progenitor marker Flk1 (also named as KDR: VEGFR2positive progenitors) (**Fig. 3g**). To evaluate TfR1^+^ cells using ferroptosis staining suitable for living single cells, we isolated E18.5 embryonic kidney TfR1^+^ cells. Approximately 36% of the isolated TfR1^+^ cells were positively stained with Mito-FerroGreen and 4-HNE (**Fig. 3h**) and 6% of the TfR1^+^ cells were triple-positive for Mito-FerroGreen, 4-HNE and TUNEL (**Fig. 3h**), indicating these cells reached the end stage of ferroptotic cell death. The percentage of 4-HNE-positive, or double 4-HNE^+^ and Liperfluo^+^ cells, in TfR1 antibody pull-downed cells, was 31% and 52%, respectively, and 8% of the TfR1-positive cells were also triple-positive for Liperfluo, 4-HNE and TUNEL (**Fig. 3h**). We used an antibody to cleaved caspase-3 to identify apoptotic cells and found a marked number of apoptotic cells in kidney tissue pre-treated with camptothecin but not in the vehicle-treated tissues (**Fig. S3f**). Further, we found no necroptotic cells in the glomeruli of the embryonic kidney, with the specificity of the necroptosis antibodies confirmed using positive control tissues and neutralized antibodies (**Fig. S3g**), suggesting that apoptosis and necroptosis may not participate in kidney morphogenesis. We also found a negative signal for FSP1 staining in the glomeruli, indicating that the FSP1 pathway does not participate in kidney ferroptosis (**Fig. S3h**).

### A distinct group of oxidized phospholipids are associated with embryonic ferroptosis

Lipid peroxidation drives ferroptosis, and we observed their presence in embryonic ferroptotic cells (**Fig. 1d, e**; **Fig 2a**). Previous findings have shown that polyunsaturated ether phospholipids (PUFA-ePLs) act as lipid peroxidation substrates leading to the induction of ferroptosis in cancer^26^. We found by lipidomic profiling that there is an accumulation of PUFA-ePLs and decreased abundance of the ester-linked PUFA chain at the sn-2 position, particularly phosphoethanolamine (PE), in TfR1^+^ cells from the embryonic kidney and lung (**Fig. 4a, b**). An enrichment analysis of the lipidomic profile also pointed to PUFA-ePLs as the dominant lipids in TfR1^+^ cells of the lung and the kidney (**Fig. S4a, b**). Among the top five PUFA-ePLs, PE(19:5e) was enriched in both kidney and lung TfR1^+^ cells (**Fig. 4a-d**). Among downregulated ester-linked PUFAs, PE(20:1_22:6) was shared by both kidney and lung TfR1^+^ cells (**Fig. 4a, b, e, f**). To determine if PUFA-ePLs are the ones oxidized, like in cancer and terminally differentiated cells^27^, we performed oxidized lipidomic profiling of TfR1^+^ and TfR1^-^ kidney cells (**Fig. 4g, h, i, j**) and found that the most abundant oxidized lipids were TG 54:8;O2 (ester), PG 43:7;O (ether), TG 52:9;O2 (ester) and PG 16-18:0;O (ester) (**Fig. 4h, i, j**). These findings indicate that a mixture of ester and ether contributes to lipid peroxidation, and a different group of increased ether and decreased ester PUFA-PLs in general lipidomic (**Fig. 4a, b, c, d, e, f**) may mainly counter lipid peroxidation in pro-ferroptotic cells. Treatment of kidney and lung TfR1^+^ cells with the ester-linked PUFA PE(16:0_22:6), which is structurally related to PE(20:1_22:6) **(Fig. 4e, f)**, significantly reduced the numbers of Mito-ferroGreen^+^TUNEL^+^ or Liperfluo^+^TUNEL^+^ cells (**Fig. 4k**), ostensibly due to the reduction of lipid peroxidation and terminal ferroptosis. The critical involvement of lipid peroxidation is further supported by the following findings. GPX4 Tg diminished whereas GPX4 cKO enhanced the presence of Mito-ferroGreen^+^4-HNE^+^TUNEL^+^ in embryonic kidney cells (**Fig. S4c**). In TfR1^-^ cells, Erastin induced the presence of Mitoferro^+^4-HNE^+^TUNEL^+^ cells (**Fig. S4d**); however, in TfR1^+^ cells, Ferrostatin-1 abrogated the presence of these triple-positive cells (**Fig. S4d**), indicating that Mitoferro positivity points to the pro- and end-stage of ferroptosis. Treatment of an oxidized PC, oxidized 1-palmitoyl-2-arachidonoyl-sn-glycero-3-phosphocholine (oxPAPC), mimicking the effect of oxidized lipids, turned TfR1^-^ kidney and lung cells into Mito-ferroGreen^+^TUNEL^+^ or Liperfluro^+^TUNEL^+^ cells (**Fig. 4l**). oxPAPC treatment did not cause Janus-like effects because it did not change the expression of Cox2 and Pdl1 (**Fig. 4e**). The above lipidomic analysis demonstrates that a group of distinct oxidized phospholipids leads to lipid peroxidation in pro-ferroptotic cells during organogenesis whereas a group of increased PUFA-ePLs arise that could possibly counteract lipid peroxidation and reverse the course of most pro-ferroptotic cells into a survival mode.

**Figure 4.**
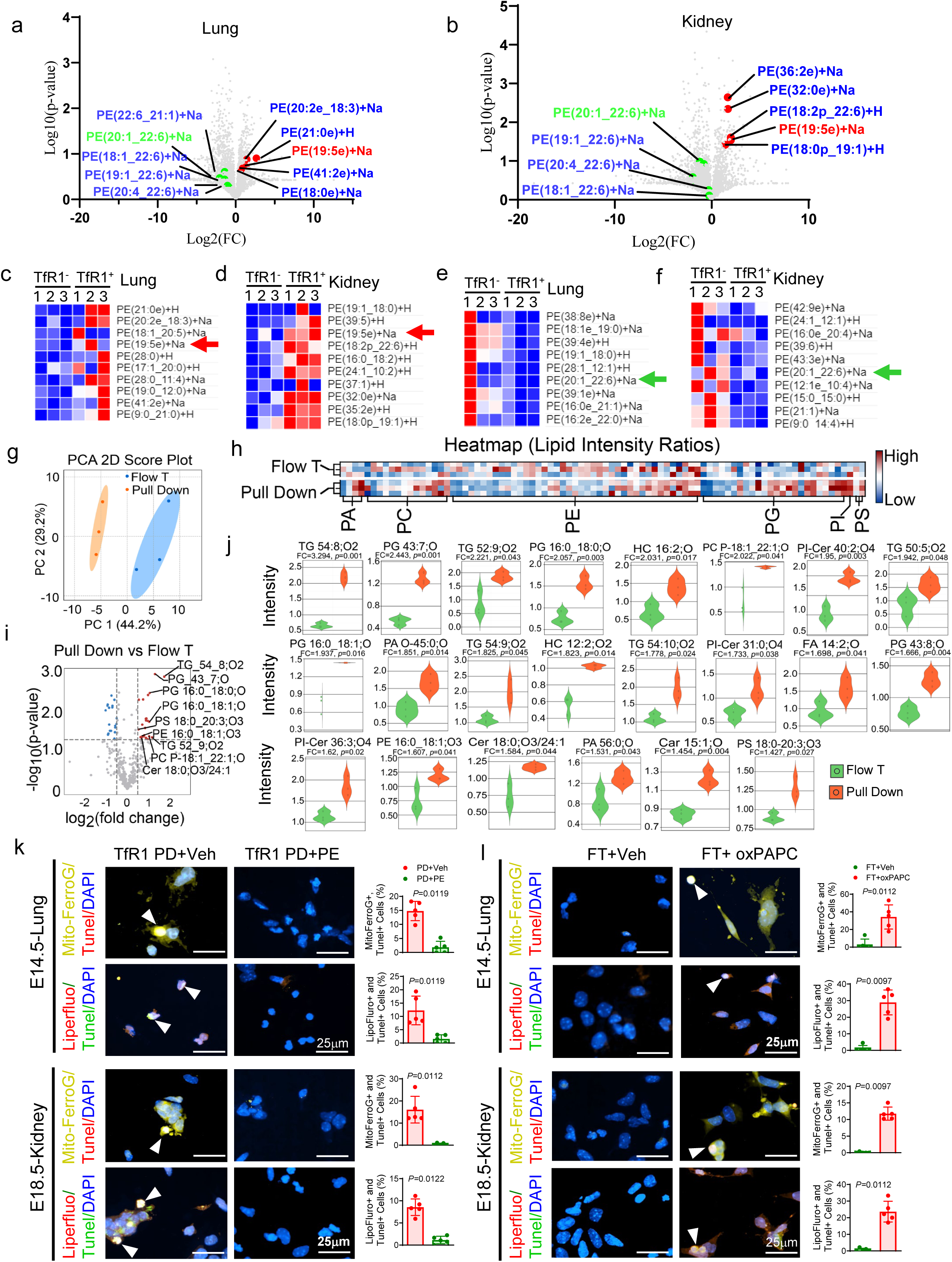
Elevation of PUFA-ePLs in TfR1^+^ cells and a mixed type of lipid peroxidation are associated with pro-ferroptotic cells. (**a**) Volcano plot of differentially detected lipids by a general lipidomics in pro-ferroptotic cells (TfR1 pull-down) compared with flow-through cells in E14.5 lung. (**b**) Volcano plot of differentially detected lipids in pro-ferroptotic TfR1 pull-down cells compared to non-ferroptotic cells (flow-through) in E18.5 kidney. Top 10 upregulated lipid in lung (**c**) and kidney (**d**), red arrows indicate the PE (19:5e)+Na was detected in both kidney and lung. Top downregulated lipid in lung (**e**) and kidney (**f**), green arrows indicate the PE (22:1_22:6)+Na was detected in both kidney and lung. (**g**) The PCA 2D Score Plot of oxidized lipidomics in TfR1 pulldown and flow-through (Flow T) cells from E18.5 kidneys. (**h**) The heatmap indicating lipid intensity ratios. Top upregulated oxidized lipids in a Volcano plot (**i**) and Violin plots for individual lipids (**j**). (**k**) Mito-FerroGreen, Lipofluro and TUNEL staining in TfR1pull-down cells from E14.5 lung and E18.5 kidney treated with PE (16:0_22:6). (**l**) Mito-FerroGreen, Lipofluro and TUNEL staining of the flow-through cells from E14.5 lung and E18.5 kidney treated with oxPAPC (oxidized 1-palmitoyl-2-arachidonoyl-sn-glycero-3-phosphocholine). Percentages of TUNEL and Mito-FerroGreen or Lipofluro positive cells in the bar graphs in **k**, **l**. FT, flow through; PD, pull down. Lung/Kidney from the embryos of five different litters (*n* = 5) were subjected for analysis and each litter consists of 8-9 embryo, and the cell samples have triplicate for each experiment. Data were analyzed by non-parametric Wilcoxon method or Mann-Whitney U test.

It has been shown that genes for peroxisomal biogenesis, including *PEX3*, *PEX10* and *PEX12,* fatty acyl-CoA reductase 1 (*FAR1*) and the endoplasmic reticulum resident enzyme 1-acylglycerol-3-phosphate *O*-acyltransferase 3 (*AGPAT3*), are required for PUFA-ePLs synthesis and increase ferroptosis susceptibility^26^. Notably, we found that all these genes were upregulated in TfR1^+^ cells (**Fig. S4f)**, suggesting that TfR1^+^ cells are primed for ferroptotic cell death.

### Single-cell RNA-sequencing identifies developmental ferroptotic cells and their signature genes

Because during induction of ferroptosis *TfR1* mRNA is not increased but its protein is translocated to the cell membrane^4^, we performed single-cell RNA-sequencing (scRNA-seq) to define the differentially-expressed genes during embryonic lung and kidney cell ferroptosis. We processed 10,000 cells from three E18.5 kidney pairs using the 10x Chromium system and 6,257 cells with an average of 980 detected genes per cell passed quality controls (**Fig. S5**). Consistent with a previous report^27^, we identified 16 embryonic kidney cell clusters, including nephron progenitors, podocytes, mesangial cells, proximal tubule cells, connecting tubule cells and glomerular endothelial cells (**Fig. S5a, b**). When examining 34 manually-curated and verified positive ferroptosis regulators, immune cells in clusters 1, 15 and 16 had the greatest number of regulators with increased expression (**Fig. S5c**). However, the expression of the major ferroptosis inhibitor Gpx4 was higher in these cells (**Fig. S5d**), indicating that immune cells do not undergo ferroptosis. As immunofluorescence detected Flk1^+^ cells (also called Kdr^+^ cells) and not all Kdr^+^ cells were ferroptotic cells (**Fig. 3f-h**), we further separated the Kdr^+^ cluster 12 into K1, K2, K3 and K4 subclusters (**Fig. S5e**). Subcluster K1 and K4 with Kdr expression had the largest number of increased ferroptosis markers (**Fig. S5f, g**). K3 had minimal Kdr expression and the highest Gpx4 expression, and K2 had the least number of positive ferroptosis markers (**Fig. S5f**). This evidence suggests that subclusters K1 and K4 cells are ferroptotic. Notably, we found that *Zeb1* expression was highly prevalent in both K1 and K4 cells, while *Alox12* expression was only prominent in K4 cells (**Fig. S5f**), indicating that these two genes may participate in kidney Flk1^+^ cell ferroptosis.

In mice, branching morphogenesis of the lung begins at E10.5 when the primary lung bud begins to invade into the surrounding mesenchyme. During this branching process, we found that Sox9 was expressed in the distal tips of the branching epithelial cells, some of which will be cleaned up by ferroptosis. Consistent with prior studies reporting Sox9 that was down-regulated at E16.5 (ref. 21), our data revealed that pro-ferroptosis begins at E10.5 in the lung primordium and reaches a peak at E14.5 and starts to decline at E16.5 (**Fig. 1b, S1a**). To further understand the role of ferroptosis in lung development, we used E14.5 whole-lung cells and TfR1 pull-downed cells to perform scRNA-seq (**Fig. 5**). In the whole lung sample, 3,916 cells with an average of 5,800 detected genes per cell were captured (**Fig. 5a**); in the TfR1 pull-downed sample, 4,762 cells with an average of 6,450 detected genes per cell were captured for downstream analysis (**Fig. 5b**). 18 cell clusters were defined in the whole lung sample with seven Sox9^+^ clusters (**Fig. 5a**). WL13 and WL14, annotated as alveolar epithelial cells (**Fig. 5a**), were the most likely Sox9^+^ ferroptotic cells because of their high TfR1 expression (**Fig. 5c, d**). The ferroptosis positive regulator Mt1 in WL14 was significantly higher than in the WL13 cluster (**Fig. 5d**). Additionally, the expression of ferroptosis negatively regulated genes, including *Noca4*, *Nfs1*, *Lpcat3*, were absent in the WL14 cluster (**Fig. 5e**), indicating WL14 is the ferroptosis cluster with high Mt1 expression. In the TfR1 pull-downed samples, 21 cell clusters with different cell type annotation were defined (**Fig. 5b**) and 13 Sox9^+^ clusters were detected (**Fig. 5f**), indicating that pro-ferroptotic cells are Sox9^+^. Because Sox9^+^ cells are epithelial cells (**Fig. 5b**), PL1-5, annotated as epithelial cells were the most likely Sox9^+^ ferroptotic cells. Even though PL3 *Mt1* mRNA level appear higher than the expression in the PL2 cluster (**Fig. 5g**), no significant difference was detected. But the ferroptosis negatively-regulated genes, including *Fancd2*, *Gclc*, *Lpcat3*, *Got1* and *Nfs1*, were not detected in the PL2 cluster (**Fig. 5h**), suggesting PL2 is the ferroptosis cluster.

**Figure 5.**
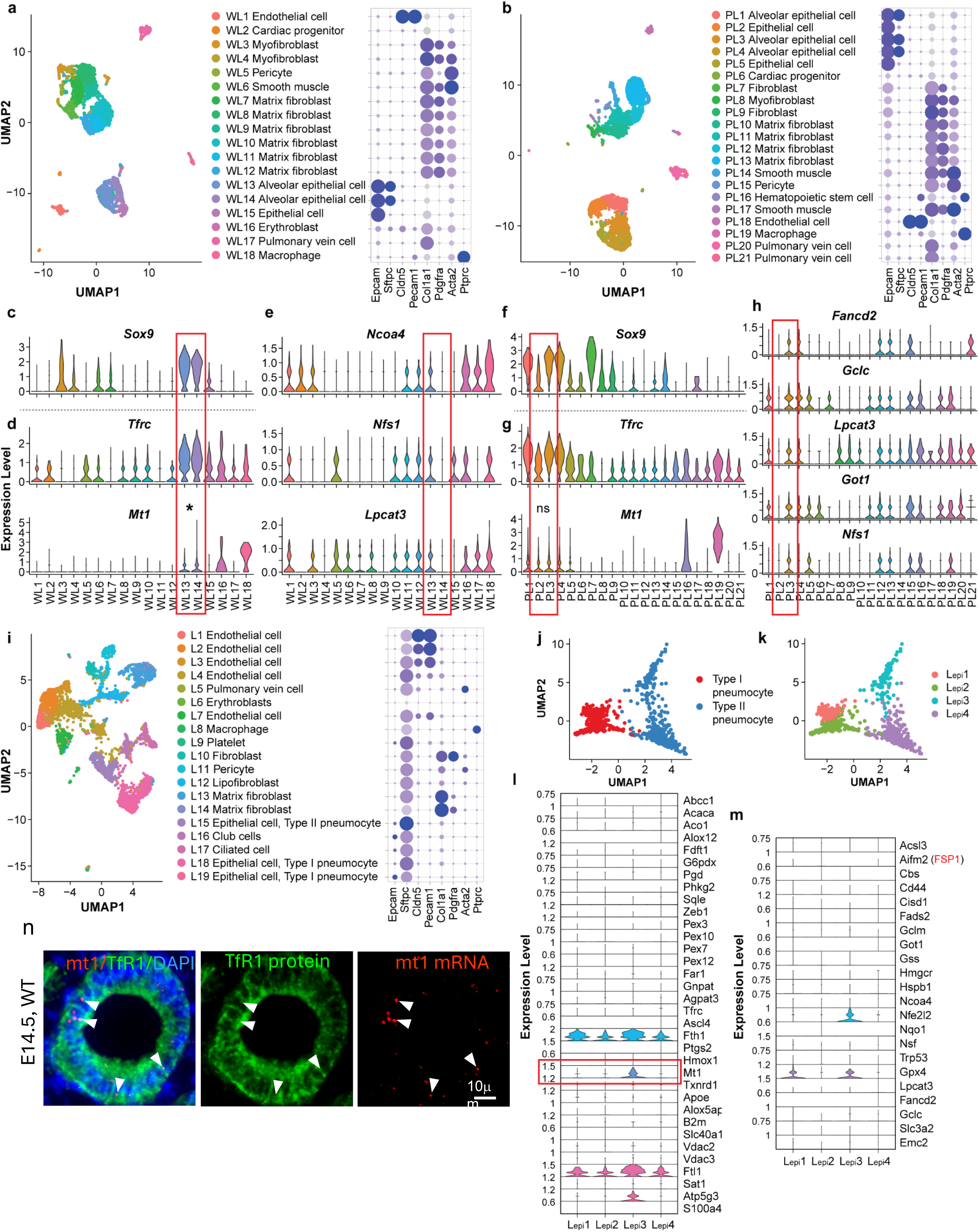
Single-cell RNA-sequencing confirms the presence of ferroptotic cells in the developing lung. (**a**) Clustering plot of E14.5 whole lung cells (*n* = 3916) by UMAP (Uniform Manifold Approximation and Projection) based on the expression of highly variable genes. (**b**) Clustering plot of E14.5 TfR1 pull down cells (*n* = 4762) by UMAP on the basis of the expression of highly variable genes. (**c**) Violin plots of Sox9 gene arranged by clusters in whole lung cells. (**d**) Violin plots of ferroptosis positive gene Tfrc and Mt1 arranged by clusters in whole lung cells; asterisk indicated significant difference compared with other clusters using Wilcoxon rank sum test. (**e**) Violin plots of ferroptosis negative gene Noca4, Nfs1, Lpcat3 arranged by clusters. (**f**) Violin plots of Sox9 gene arranged by clusters in TfR1 pull down cells. (**g**) Violin plots of ferroptosis positive gene Tfrc and Mt1 arranged by clusters in TfR1 pull down cells; no significant difference was detected in these two clusters using Wilcoxon rank sum test. (**h**) Violin plots of ferroptosis negative gene Fancd2, Gclc, Got1, Nfs1 arranged by clusters. (**i**) Clustering plot of E16.5 whole lung cells (*n* =6147) by UMAP based on the expression of highly variable genes. (**j**) Sub-clustering epithelial cells into type I and type II pneumocyte. (**k**) Trajectory analysis of four epithelial subclusters. (**l**) Violin plots of ferroptosis positive gene arranged by clusters in E16.5 whole lung cells. (**m**) Violin plots of ferroptosis negative gene arranged by clusters in E16.5 whole lung cells. (**n**) RNAscope dual staining (Mt1 RNA red; TfR1 green) in E14.5 lung distal epithelium. WL, E14.5 whole lung; PL, E14.5 pull down lung; L, E16.5 whole lung.

Next, we aimed to determine whether ferroptotic cells with the same gene signature can be detected at a later stage of lung development. E16.5 is a timepoint when alveolarization begins. So next we used whole-lung samples at E16.5 for scRNA-seq and annotated 19 cell clusters (**Fig. 5i**). Because ferroptotic cells were detected in the distal epithelium airways (**Fig. 2a**), which are composed of type I and type II pneumocytes at E16.5, clusters 15, 18 and 19 containing these cells were sub-clustered into four sub-clusters L1, L2, L3 and L4 (**Fig. 5j, k**). Among these four sub-clusters, L3 had the highest number of positive ferroptosis markers, with the two genes *Mt1* and *Atp5g3* as its unique markers (**Fig. 5l**). Although the major ferroptosis inhibitor GPX4 was expressed in the L3 cluster (**Fig. 5m**), *Mt1* mRNA overlapped with TfR1 staining signals in the distal airway epithelium (**Fig. 5n**), supporting L3 cells with the unique marker Mt1 as ferroptotic cells in the lung.

### Loss or overexpression of *Gpx4* alters lung distal airway morphogenesis

There are two major pathways that inhibit ferroptosis: the GPX4-GSH pathway and the ferroptosis suppressor protein 1 (FSP1)-coenzyme Q10 (CoQ) pathway^16,28^. It appears that FSP1 (originally named as Aifm2) is not critically involved in embryonic cell ferroptosis as it was not significantly changed in our scRNA-seq analyses (**Fig. 5m, S5d, g**). On the other hand, GPX4 showed a marked difference between non-ferroptotic and ferroptotic cells (**Fig. 5m, S5g**), indicating GPX4 regulates ferroptosis during embryonic development.

Next, we used loss- and gain-of-function approaches to assess the impact of different GPX4 levels on lung development. Conditional *Gpx4* deletion in all lung lineage cells using *Nkx2.1* promoter-driven Cre mice crossed with *Gpx4*^flox/flox^ mice resulted in an increase in the number of TfR1^+^4-HNE^+^TUNEL^+^ triple-positive cells in the lung (**Fig. 2a**) and an significantly increased abundance of MDA (Malondialdehyde), another lipid peroxidation marker, in embryonic lungs (**Fig. S6a**), as well as cyst-like structures at the distal epithelial tips leading to large airspaces and disrupted septae organization inside the distal airways (**Fig. 6a right panel, Fig. S6b**). We divided the lung with different degrees of cyst burden into minor (< 10% area), medium (10-20% area) and severe (> 20% area) and found that the conditional *Gpx4* deletion displayed different degrees of cyst burden (**Fig. 6b, Fig. S6b**).

**Figure 6.**
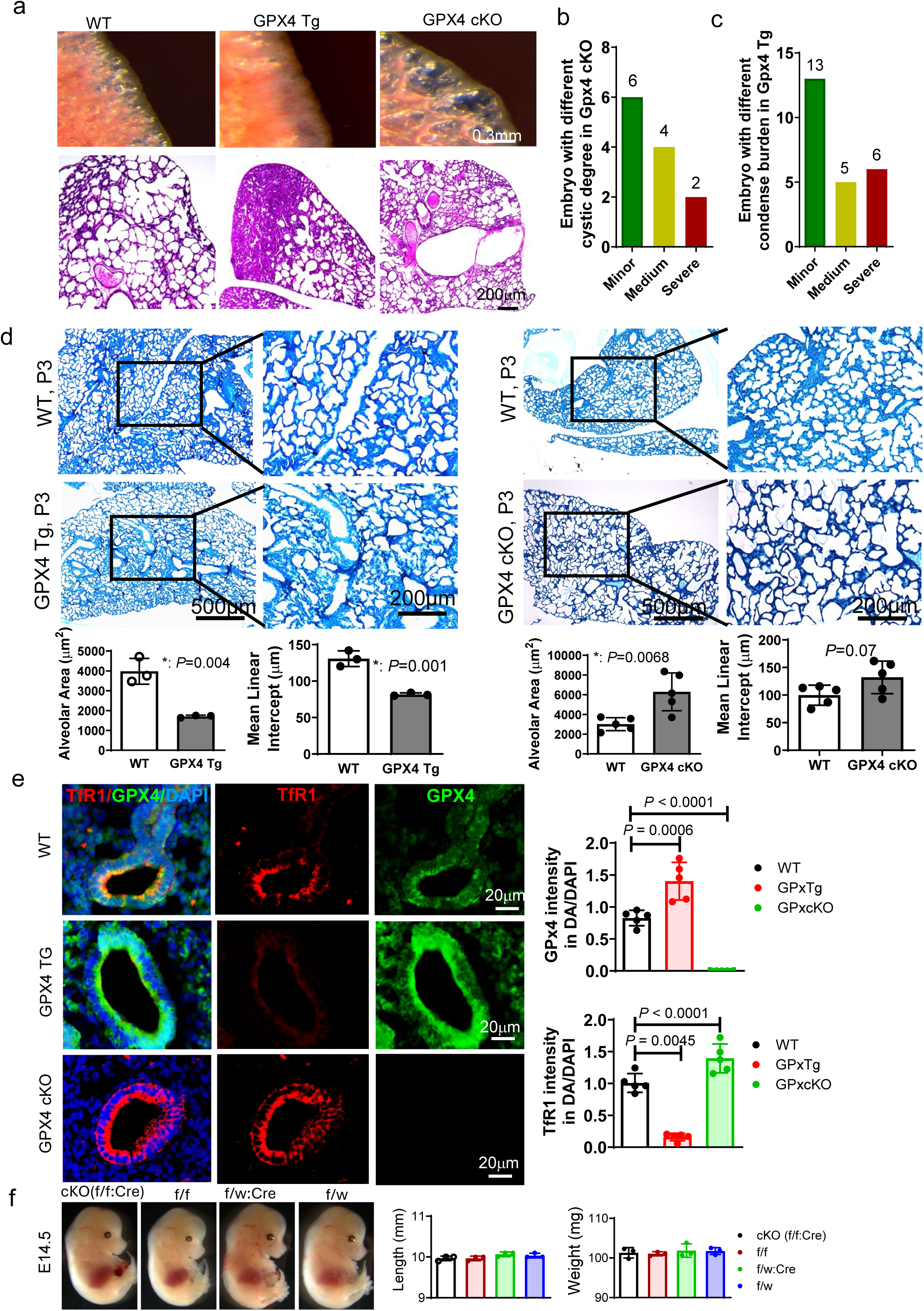
*Gpx4* knockout or overexpression leads to abnormal lung development. (**a**) Top panel, gloss morphology of P3 lung in WT, *Gpx4* TG and *Gpx4* cKO groups; middle, higher magnification imaging indicates different alveoli morphology in different groups; bottom, H&E staining shows the abnormal lung structures in *Gpx4* TG and *Gpx4* cKO groups. (**b**) Number of embryos with different cystic degrees in *Gpx4* cKO mouse; minor, < 5%/section; medium, 5-10%/section; severe, > 10%/section. Quantification of septal thickness and (**c**) Number of embryos with different condensed burden in *Gpx4* TG mouse; minor, < 10%/section; medium, 10-20%/section; severe, > 20%/section. (**d**) Lung stereology measurement for alveolar area and mean linear intercept from 5 embryos per group. *P* values from Student’s t test. (**e**)TfR1 and GPX4 immunostaining for E14.5 lungs from WT, *Gpx4* TG and *Gpx4* cKO mice. Percentages of the TfR1, GPX4 or double TfR1 and GPX4 positive distal epithelial cells from 5 lungs (embryos) are presented in graphs. *P* values from one-way ANOVA followed by *Tukey* test. (**f**) E14.5 embryos from GPX4 cKO and their control and the measurement of embryonic length and weight from 3 embryos per group. One-way ANOVA followed by *Tukey* test.

Transgenic overexpression of *Gpx4* diminished the occurrence of TfR1^+^4-HNE^+^TUNEL^+^ triple-positive cells in the lung (**Fig. 2a**), decreased MDA expression (**Fig. S6a**) and led to alveolar collapse (**Fig. 6a middle panel**). Additionally, *Gpx4* overexpression resulted in extensive areas of large dense patches with minimal airspaces (**Fig. 6a, c, Fig. S6c, d**)..*Gpx4* transgenic overexpression decreased the alveolar area and the mean linear intercept in embryonic lungs (**Fig. 6d**). In contrast, *Gpx4* deletion caused increases in alveolar area and mean linear intercept in embryonic lungs (**Fig. 6d**). *Gpx4* transgenic overexpression diminished TfR1 protein (**Fig. 6e**), MT1 protein and mRNA expression (**Fig. S6e, f**) but did not affect overall embryonic development at E18.5 (**Fig. S6g**). GPX4 deficiency in the lung increased TfR1 expression in the distal epithelium (**Fig. 6e**), increased the mRNA and protein expression of Mt1 (**Fig. S6e, f**), but did not affect overall embryonic development at E14.5 (**Fig. 6f**). Thus, GPX4-regulated epithelial cell ferroptosis is crucial for proper distal epithelial branching morphogenesis of the lung.

### *Gpx4* deletion or transgenic overexpression causes defective renal morphogenesis

To assess the role of GPX4 in Flk1^+^ cell ferroptosis and its effects on kidney development, we used the *Flk1* promoter-driven Cre mice crossed with the *Gpx4*^flox/flox^ mice to delete the *Gpx4* gene specifically in Flk1^+^ cells. *Gpx4* deletion in Flk1^+^ cells resulted in a greater number of ferroptotic TfR1^+^ cells inside the glomeruli (**Fig. 1e, 2b**) and led to duplex kidney formation in (1/16) 6.3% of the kidneys and (6/16) 37.5% of hydroureter (**Fig. 7a, b, Fig. S7a**).

**Figure 7.**
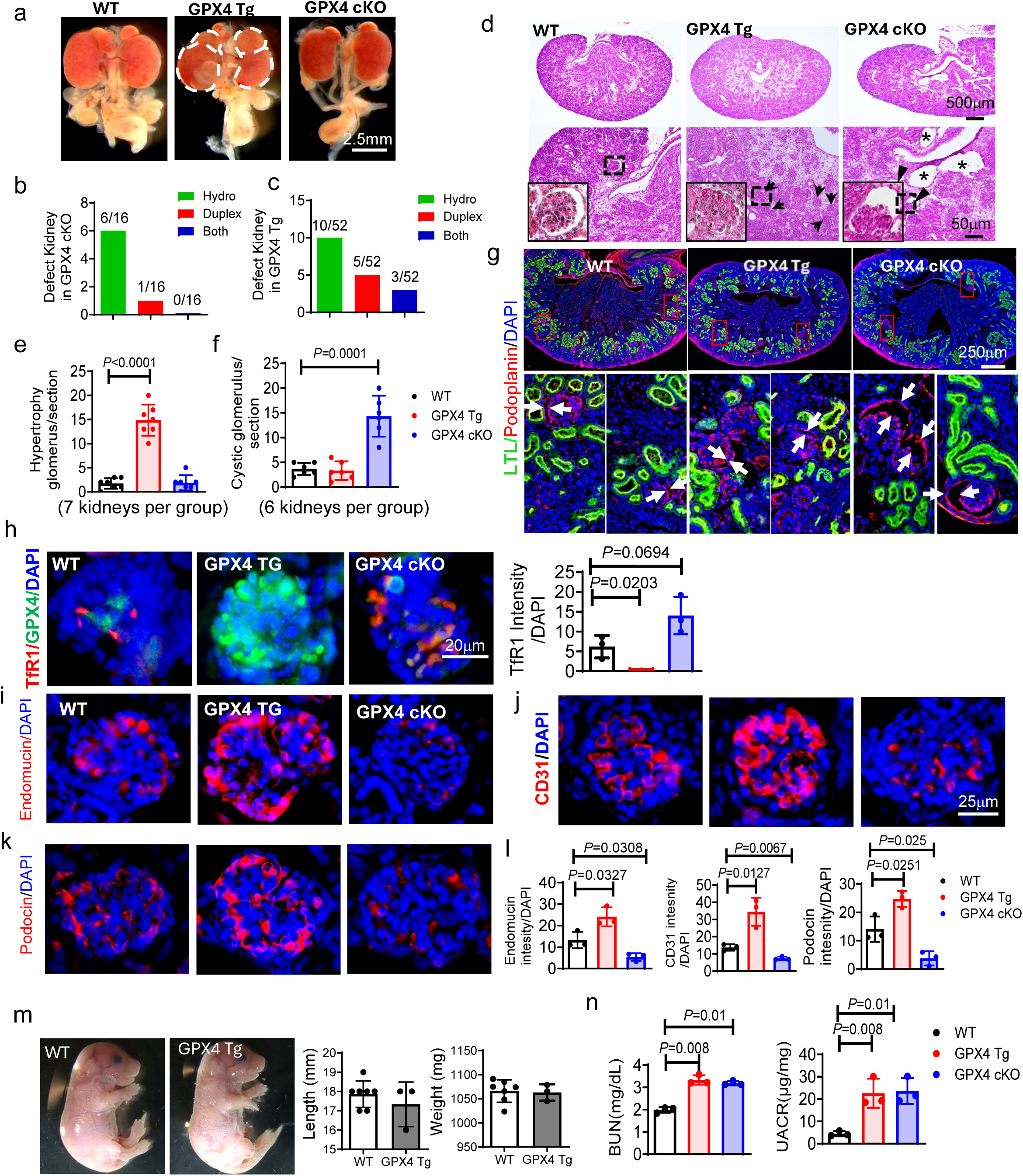
*Gpx4* deletion or overexpression interferes with proper kidney development. (**a**) Gloss morphology of E18.5 kidney in WT, *Gpx4* TG and *Gpx4* cKO groups; the dotted lines indicate the duplex kidney. Kidney with defects including hydroureteronephrosis/hydronephrosis, duplex kidney, hydroureteronephrosis/hydronephrosis plus duplex kidney in *Gpx4* cKO (**b**) and *Gpx4* TG (**c**). (**d**) H&E staining of E18.5 kidney in WT, *Gpx4* TG and *Gpx4* cKO groups; asterisk indicated cystic kidney; black arrows indicated the hypertrophy glomeruli; black arrowheads indicated the glomeruli with enlarged Bowman’s space. Corresponding quantification of hypertrophic (**e**) cystic (**f**) glomerulus in WT, *Gpx4* TG and *Gpx4* cKO groups. (**g**) Representative images of LTL (Lotus Tetragonolobus Lectin) and Podoplanin double staining to mark the proximal tube and renal corpuscles in WT, *Gpx4* TG and *Gpx4* cKO groups. Arrows point to the Bowman’s spaces. Representative images of TfR1 and GPx4 immunostaining (**h**) Endomucin immunostaining (**i**) CD31 (**j**) and podocin (**k**) immunostaining of E18.5 kidney in WT, *Gpx4* TG and *Gpx4* cKO groups; corresponding quantification was presented in (**l**). E18.5 embryos from GPX4 Tg and WT (**m**). *n* = 3 embryos for *Gpx4* Tg and *n* = 7 for WT. Renal functions determined by blood urea nitrogen (BUN) and urine albumin/creatinine ratio (UACR) of pups 2 weeks after birth (**n**). Three embryos from each litter and four to five sections per embryo were stained, and an average for signal intensity was obtained for that litter. Hydro, hydroureteronephrosis/hydronephrosis; duplex, duplex kidney; both, hydroureteronephrosis/hydronephrosis plus duplex kidney. *P* values from One-way ANOVA followed by *Tukey* test (**e, f**).

In the gain-of-function approach, we found that transgenic *GPX4* overexpression resulted in a lower number of glomerular TfR1^+^ ferroptotic cells (**Fig. 1e, 2b**), an induction of (5/52) 9.6% duplex kidneys, (10/52) 19.2% with hydroureter and (3/52) 5.8% duplex kidneys plus hydroureter (**Fig. 7a, c, Fig. S7b**) and dense renal corpuscles with a lack of Bowman’s spaces (**Fig. 7d, e, f, g**). All kidneys with *Gpx4* deletion in Flk1^+^ cells exhibited enlarged glomerular Bowman’s spaces with shrinkage of renal corpuscles, leading to cyst-like structures in the kidneys (**Fig. 7d, e, f, g**).

Flk1^+^ cells are the progenitors of endothelial cells. Thus, to determine if endothelial cell differentiation was affected by *Gpx4* deletion in Flk1^+^ cells, we used the endothelial cell marker CD31, as well as the glomerular endothelial marker endomucin, for immunostaining. We found that while *Gpx4* deletion and transgenic *Gpx4* overexpression increased and decreased TfR1^+^ cells in the glomeruli, respectively, (**Fig. 7h**), both endomucin and CD31 (**Fig. 7i, j**) expression were significantly lower in the *Gpx4* knockout kidney compared to those of wild-type kidneys, leading to a reduced filtration area in the glomerular (**Fig. 7k, l**). Conversely, we found elevated endomucin (**Fig. 7i, l**) and CD31 (**Fig. 7j, l**) expression in the *Gpx4* transgenic kidneys, resulting in a disorganized glomerular filtration area, labeled by podocin (**Fig. k, l**). The impact of *Gpx4* deletion or overexpression on ferroptosis was further confirmed by increased or decreased MDA abundance in the embryonic kidney, respectively (**Fig. S7c**), as well as protein and mRNA levels of the Flk1^+^ cell ferroptosis signature gene *Alox12* (**Fig. S7d, e**). Although either *Gpx4* deletion or overexpression did not affect overall embryonic development (**Fig. 7m, S7f**), physiologically we found that renal function 2 weeks after birth, as determined by urine albumin/creatinine ratio (UACR) and blood urea nitrogen (BUN), showed impairment in both the conditional deletion and transgenic *Gpx4* overexpressing mice (**Fig. 7n**). Thus, glomerular Flk1^+^ cell ferroptosis is required for proper kidney development.

## Discussion

### The identity of ferroptotic cells in development

Under pathological conditions, terminally-differentiated epithelial cells in the adult lung and kidney display ferroptosis^29,30^. Concomitant iron accumulation and lipid peroxidation are key features of pathological ferroptosis^29^. But a role for ferroptosis in embryonic organ development has been poorly studied, especially during the mid-to late-stages of gestation. Here, we detected ferroptotic Sox9^+^ epithelial progenitors in the developing lung and ferroptotic Flk1^+^ cells in the embryonic kidney with triple positivity for TfR1, 4-HNE and TUNEL during normal mouse development. These embryonic ferroptotic cells share the key signatures of iron accumulation and lipid peroxidation with adult ferroptotic cells as both express TfR1 and have elevated levels of the lipid peroxidation marker 4-HNE. In the lung, it appears that iron accumulation primes the epithelial progenitors towards ferroptosis because all distal airway progenitors are TfR1-positive but only 23.4% of these TfR1^+^ cells in the distal airway lumen are also positive for 4-HNE and TUNEL. Likewise, only 32.3% of the TfR1^+^ cells are positive for 4-HNE and TUNEL in the kidney, indicating that iron accumulation precedes lipid peroxidation. While it is infeasible to trace down the double TfR1- and 4-HNE-positive cells *in vivo*, reliable *in vitro* ferroptosis assays detected similar percentages of ferroptotic cells (double positive for 4-HNE and TUNEL) in TfR1^+^ cells extracted from the lung or the kidney, as those for double TfR1- and 4-HNE-positive cells *in vivo*.

TfR1 antibodies allow the detection of ferroptotic cells in cultured cells^4^. In tumor sections, TfR1^+^ cells overlapped with MDA^+^ cells^4^. In embryonic tissues, we found that only a fraction of TfR1^+^ cells also exhibit lipid peroxidation. Thus, double TfR1 and 4-HNE staining is more reliable than single staining for the detection of embryonic cell pro-ferroptosis. TUNEL staining identifies a portion of TfR1^+^ and 4-HNE^+^ cells that undergo cell death. In the distal lung epithelium where TfR1^+^ and 4-HNE^+^ cells exist, no necroptosis are detected. On the other hand, apoptotic cells in the developing lung exclusively occur in the mesenchyme surrounding epithelial airway buds, providing space for outgrowth of the lung bud^31^.

In the kidney, we found that Flk1^+^ progenitors undergo ferroptosis in the glomeruli, but not in the S-shaped body, the precursor of the glomeruli. Flk1^+^ progenitors differentiate into endothelial cells that line the glomerular capillaries and form the glomerular basement membrane ^32^. We hypothesize that ferroptosis in these cells limits the overproduction of endothelial cells inside the glomeruli to ensure proper capillary formation. In the lung, we found that the ferroptotic cells are Sox9^+^ cells, which reside in the distal airway tips^21^. These cells eventually differentiate into type 1 and type 2 pneumocytes^21^. We hypothesize that during lung morphogenesis ferroptosis is a mechanism to eliminate excessive Sox9^+^ cells, whereas it is apoptosis in mesenchymal cells, not epithelial cells, that critically participates in branching morphogenesis during lung development^33^.

### Lipid peroxidation of mixed types of lipids and the counteractive effect of ether PUFA in development ferroptosis

Either saturated or unsaturated ether glycerophospholipids are the known substrates for lipid peroxidation that primes cells for ferroptosis^26,34^. However, we found by lipidomic profiling that a group of mixed lipids are oxidized but not particularly ether PUFAs, which are elevated in pro-ferroptotic TfR1^+^ cells as determined by general lipidomics. Because most TfR1^+^ cells do not eventually commit ferroptosis, there must be a counteractive mechanism to halt the progression of these cells from pro-ferroptotic to ferroptotic. Ether PUFAs can act as reducing agents^35^ and thus increased levels of these lipids may prevent the majority of the TfR1^+^ cells from undergoing ferroptosis. Our further evidence showed that treatment with the ester-linked PUFA PE(19:1_22:6) prevents TfR1^+^ lung and kidney cells from undergoing ferroptosis, and treatment of an oxidized phospholipid oxPAPC induces ferroptosis in TfR1^-^ cells, supporting the notion that lipid peroxidation is a key driver of ferroptosis during organ morphogenesis.

Peroxisome enzymes are required for ether PUFA production^26,34^. Indeed, peroxisome enzymes, including PEX3, PEX10, PEX12, FAR1 and AGPAT3, are essential for ether lipid biosynthesis, and we found that their levels are increased in pro-ferroptotic cells during development. Besides the ROS generated by Gpx4 downregulation in development, lipoxygenases can also oxidize lipids as Alox12 has been identified as a contributor to lipid peroxidation in the kidney and Alox12 induces lipid peroxidation by stimulating lipoxygenase expression during ferroptosis^36,37^. Thus, our data suggests that decreased Gpx4-induced ROS and increased lipoxygenase expression contribute to lipid peroxidation in Sox9^+^ lung cells and Flk-1^+^ kidney cells during organ morphogenesis.

### Iron accumulation in embryonic ferroptotic cells

TfR1 is the major receptor to take up ferric-bound transferrin. Consistent with previous findings^4^, we found that *TfR1* mRNA is not upregulated during ferroptosis. TfR1 is constantly recycled back to the cell membrane after its delivery of the ferric-bound transferrin to the endosome^38^. It appears that the TfR1 positivity in Sox9^+^ lung and Flk1^+^ kidney cells are due to enhanced TfR1 membrane translocation. The observed increase of intracellular ferric in TfR1^+^ cells depends on membrane localized TfR1. Lipid peroxidation occurs in a portion of the TfR1^+^ cells, suggesting that TfR1 primes the cells for ferroptosis, but lipid peroxidation executes ferroptosis only in TfR1^+^ cells. Only a portion of TfR1-positive cells are primed to undergo ferroptosis, indicating that iron enrichment is indispensable for distal airway epithelial cell function. Iron is important for fetal lung development and iron-deficient pregnancy results in offspring with lung diseases, such as inflammation^39^.

### Ferroptosis in lung branch morphogenesis

Sox9 is exclusively expressed in the distal lung epithelial cells before E16.5 and its expression decreases afterwards^21^. Sox9^+^ cells subsequently differentiate into type 1 and type 2 alveolar cells^21^, and these cells are crucial for postnatal alveolarization. Selective Sox9^+^ cell ferroptosis may restrict the number of type 1 and type 2 alveolar cells and, thus, ensure the proper size of the airspace during development. *Gpx4* Tg expression blocks Sox9^+^ cell ferroptosis and results in a collapsed airway space. Although *Gpx4* Tg expression is not specific to Sox9^+^ cells, it inhibits Sox9^+^ cells because only these cells are primed for ferroptosis in the developing lung. Conversely, conditional *Gpx4* deletion using the lung early progenitor marker *Nxk2.1*-driven Cre increases the rate of ferroptosis only in distal epithelial airway cells and causes dilated airspaces in the developing lung, suggesting that the distal epithelial airway cells are primed by accumulated iron for ferroptosis and *Gpx4* deletion enhances lipid peroxidation in these cells leading to increased rate of ferroptosis.

### Ferroptosis in glomerular capillary looping

At E12.5, the developing mouse renal blastema has its own pool of Flk1^+^ endothelial progenitor cells^32^. Because ferroptosis in Flk1^+^ progenitors is not present in other parts of the embryo, Flk1^+^ progenitors in the glomeruli are a unique population of cells that acquire susceptibility to ferroptosis. Flk1^+^ progenitors surrounding the comma-shaped body and “streaming” into the vascular cleft of the S-shaped body^40^, precursors of the glomerulus, do not show signs of ferroptosis, indicating that ferroptosis in Flk1^+^ progenitors occurs in a spatial-temporal fashion and only inside the developing glomerulus. During glomerular capillary looping and maturation, superfluous Flk1^+^ progenitors are removed by ferroptosis, and most of these Flk1^+^ progenitors have been already differentiated into endothelial cells that form the capillary tuft in the Bowman’s space with multiple and parallel branches^40^. On the other hand, excessive endothelial cells in the glomerulus are removed after birth via apoptosis, which is required for capillary lumenization^41^. We found here that enhancing Flk1^+^ progenitor ferroptosis via *Gpx4* deletion resulted in few and distended capillary loops in the glomerulus and the formation of “cyst-like” glomeruli with enlarged Bowman’s space. Preventing Flk1^+^ progenitor ferroptosis through *Gpx4* Tg expression resulted in “overcrowded” and disorganized capillary microvessels in the glomerulus with a loss of the Bowman’s space. Thus, Gpx4-regulated Flk1^+^ progenitor ferroptosis is indispensable for proper glomerular capillary looping.

In summary, we identify ferroptotic cells during major organ morphogenesis. Ferroptosis during development occurs in selective cell-types in different organs and in a spatial-temporal manner. Iron accumulation with concomitant lipid peroxidation causes ferroptosis. Gpx4 is a major regulator for developmental ferroptosis and gain-or loss-of-function approaches that deplete or enhance ferroptosis lead to defects in lung alveolarization and kidney glomerular capillary looping. These findings support the notion that ferroptosis acts embryonically to remove superfluous cells during morphogenesis to ensure proper organ development.

## Methods

### Animals work

All animal procedures were approved by the Institutional Animal Care and Use Committee of our University. All the mice were on the C57/BL6 genetic background. The mice were housed at 20-22°C under a 12 h light/12 h dark cycle with free access to standard diet and water. Male and female pairs were set up at 3:00 p.m and an E0.5 pregnancy was established when a vaginal plug was identified in the morning of the following day. Embryos were harvested at 2:00 p.m for subsequent experiments at different developmental stages. *Gpx4* transgenic male mice were mated with WT female to generate WT and *Gpx4* TG embryos. *Nkx2.1*-Cre (Stock No. 008661), and *Flk1*-Cre mice (Stock No. 018976) were purchased from the Jackson Laboratory. Mice with a targeted deletion of *Gpx4* in lung (*Nkx2.1*-Cre; *Gpx4*^loxp/loxp^) and kidney (*Flk1*-Cre; *Gpx4*^loxp/loxp^) were yielded by crossing the *Gpx4*-floxed mice with *Nkx2.1*-Cre, and *Flk1*-Cre mice, respectively. Cre-negative littermates were used as control throughout the experiments.

### Primary cells isolation and culture

Primary lung and kidney cells were isolated from fresh tissues. Isolation method will be described in following single cell dissociation. The cells were maintained in Endothelial Cell Growth Basal Medium-2 (EBM-2) supplemented with 10% FBS at 37°C in a humidified atmosphere of 5% CO_2_ in incubator. The cells were ready for downstream experiments after they attached to the plate. PE (16:0_22:6) and oxPAPC were added to the medium in a 1 μm and 30 μg/μl concentration respectively when lipid treatment was carried out.

### Animal Tissue collection, processing and hematoxylin-eosin staining

Fresh tissues were fixed using Carnoy’s fixative and fully dehydrated and paraffin embedded and cut into 5-10 μm sections. For frozen section preparation tissue was fixed in 4%-paraformaldehyde/PBS solution followed by dehydrated with 10-30% sucrose solution. Cryo blocks were prepared in OCT compound and cut into 10 μm sections. Hematoxylin-eosin (H&E) staining was performed using 5 μm thick paraffin embedded sections after deparaffinization of the tissue using graded xylene and alcohol. Staining was performed using modified Harris hematoxylin (1:1 in water) followed by treatment with lithium carbonate. After extensive wash tissue was treated with Eosin Y, further washed and passed through graded alcohol and xylene for mounting.

### Immunofluorescence, Immunocytochemistry and TUNEL Staining

For Immunofluorescence both paraffin embedded and frozen sections were used and were performed following the protocol used previously^42^. For paraffin embedded tissue deparaffinization was performed before antigen retrieval. Air dried frozen sections directly emerged in antigen retrieval solution. Antigen retrieval was performed by autoclaving at 120°C for 10 min in citrate buffer. After blocking and permeabilization with 10% normal goat serum in 0.3% saponin PBS for 1 h at room temperature. The embryonic kidney sections were further blocked with Fc (TruStain FcX™ PLUS, biolegend) to reduce the background effect of immunofluorescence. Slices were incubated with primary antibody diluted in antibody dilution solution containing 0.1% BSA,0.1% Gelatin in 0.05% Tween20/PBS at 4°C overnight. Details about the antibody are presented in the supplementary information. For visualization sample were incubating with Alexa flour labelled specific secondary antibody (F ab fraction) 1 h at 37°C temperature, followed by nuclei counter-stained with DAPI. For Immunocytochemistry 4%-paraformaldehyde/PBS fixed cells were treated with 0.2% TritonX or Saponin/PBS for permeabilization. Following blocking with 5% BSA cells were incubated with primary antibody diluted in antibody dilution solution at 4°C overnight. Microscopy was performed using Nikon CSU-W1 Spinning Disk Field Scanning Confocal microscope and Nikon Light microscope. Images were captured and processed using NIS element software.

Tunel staining was performed using In Situ Cell Death Detection Kit (TMR red or Fluorescein, Roche and ApopTag Fluorescein Direct In Situ Apoptosis detection kit, Sigma) following the manufacturer’s instructions. Briefly, after incubation with secondary antibody, the working strength enzyme and substrate mix was applied to the 20-30 μm thick slices or cell samples and incubated at 37°C for 1 hr. The nuclei were counter-stained with DAPI for imaging. Serial sections from whole lungs or kidney were subjected to microscopy for the detection of Tunel positive cells.

### Iron Staining (Prussian Blue Stain) and confirmation of TfR1 positive cells

Paraffin embedded tissue section was subjected for immunofluorescence and Prussian blue staining (Perl) to detect iron deposits in Ferroptotic cells. After deparaffinization tissue sections were first subjected for TfR1 immunostaining and mounted with 50% glycerol for microscopy. After documentation of the TfR1 microscopy, cover glass was removed by keeping in cold PBS. Following that, Perl staining was conducted according to the manufacturer’s protocol (AbCam). Briefly, working iron staining solution was prepared by mixing equal volume of Potassium Ferrocyanide Solution and Hydrochloric Acid Solution and then applied to the slices for 5-10 minutes at room temperature. Nuclei were labeled with Fast Red Solution for 5 minutes. The ionic iron can be visualized by reacting with acid ferrocyanide to produce a blue color. Microscopy was performed using Nikon Eclipse Ni microscope and images were captured and processed using NIS element software.

### Live and dead cell staining

Direct detection of the live and dead cells in the embryonic lung and kidney tissue was performed by using membrane specific dye MemBrite Fix Cell Surface Staining Kits (biotium) and Nuclear stain SYTOX Orange (live and dead cell staining). A mixture of these two dyes (1:1000 dilution in nuclease& protease free PBS) was applied to embryos through whole animal perfusion after washing the whole-body blood with 10 ml ice cold PBS. 5ml of the dye solution were used for staining to pass the body by 50 minutes. Immediately after perfusion, lungs and kidney were collected, washed, and fixed using 4% PFA for 3 hours (light protected). Pre-stained fixed lung and kidneys tissue then used to prepare agarose block using Low Melting Point Noble Agar. 50 μm thick floating sections were prepared using vibratome (Leica VT1000S) and incubated with DAPI for nuclear staining. After an extensive wash with PBS thick sections were mounted and subjected for microscopy using Nikon CSU-W1 Spinning Disk Field Scanning Confocal microscope.

### Electron microscopy, tissue and cell processing

For lung and kidney tissue preparation for electron microscopy, embryos were perfused and fixed transcardially using 2% Paraformaldehyde, 2.5% Glutaraldehyde in 0.1 M Sodium Cacodylate buffer pH 7.4. After collecting the E14.5 lung or E18.5 kidney from the embryo, samples were further kept in fixative for 1hr at room temperature followed overnight at 4°C overnight. Fixed lungs and kidneys further subjected to agarose block preparation using Low Melting Point Noble Agar and 100 μm thick floating sections were prepared using vibratome (Leica VT1000S). Section were placed in 2% Paraformaldehyde, 2.5% Glutaraldehyde in 0.1 M Sodium Cacodylate buffer pH 7.4 for overnight at 4°C. Following day aldehydes were washed and quenched with 100 mM Glycine in Cacodylate buffer, and post fix with 1% OsO4. Following an extensive dehydration of the tissue with graded alcohol the tissue blocks were transferred to acetone. The tissues were infiltrated and embedded in Epon. Ultrathin sections of 200 nm are cut and mounted on a silicon wafer. Images were taken by using Apreo 2 SEM Scanning electron microscope (UMB electron microscopy core).

For cells, after fixation with 2% Paraformaldehyde, 2.5% Glutaraldehyde in 0.1 M Sodium Cacodylate, the cells were embedded in 4% agarose. Following OsO4 post fixation, dehydration and embedding were carried out as described earlier ^43^. 50 nm ultra-thin sections were cut and subjected to imaging using Tecnai T-12 transmission electron microscopy (TEM).

### RNAscope Multiplex Fluorescent Assay combined with Immunofluorescence

RNAscope fluorescent assay combined with immunofluorescence staining was carried out following manufacturer’s protocol. Briefly, tissue sections were deparaffinized by running the slices through xylene and 100% ethanol. Then the RNAscope hydrogen peroxide was applied to the slices for 10 minutes at room temperature. Target retrieval was performed by submerging the slices into the mildly boiling target retrieval reagents solution for 15 minutes. After that, RNA probes were added to cover the slices and incubated for 2 hours at 40°C. AMP1, AMP2, AMP3 were applied to the slices in sequence after the probe hybridization. After washing, HRP-C1 was added to each slice and incubated for 15 minutes at 40°C, then the diluted Opal^TM^ 520 was applied to each slice and incubated for half hour at 40°C. HRP blocker was added to the slices, and PBS washing was conducted. Then the immunofluorescence procedures, including serum blocking, primary antibody incubation and specific secondary antibody incubation, were carried out in sequence using TBS buffer. Finally, the nuclei were counter-stained with DAPI for imaging.

### Mito-FerroGreen/Liperfluo Live Cell Staining coupled immunostaining

According to the manufacturer’s instructions, cells were washed with PBS twice before they were cultured in serum-free HBSS medium with 5 μmol/L of Mito-FerroGreen or 1 μmol/L of Liperfluo working solution for 30 minutes at 37°C in a 5% CO2 incubator. Following the incubation cells were washed and fixed with 4% PFA and subjected for immunostaining with specific antibodies as described earlier. Cells were visualized under a Nikon CSU-W1 Spinning Disk Field Scanning Confocal microscope (Nikon, Japan).

### Magnetic Microbeads-based Live Cell Pull-down Assay

Lungs collected from E14.5 embryos were chopped and incubated with enzyme mix (Lung Dissociation Kit; Miltenyi Biotec) for 20 minutes at 37°C water bath under gently rotation every 5 minutes. After digestion, cells were filtered by 70 μm strainer and washed with serum containing medium to stop further digestion.

Kidneys obtained from E18.5 embryos were minced and incubated with collagen IV for 10 minutes at 37°C water bath under gently rotation every 5 minutes. The following procedures were the same as for the lung.

Magnetic microbeads-based live cell pull-down assays were conducted following the manufacturer’s instruction. After removal of the red blood cells, cells were washed with PBS and centrifugation at 300×g for 5 minutes. The ells were resuspended with MACS buffer (1% FBS+2Mm EDTA in PBS) and cultured with CD71(TfR1)/CD309 (Flk1) or E-cadherin antibody for 10 minutes at 4°C while protected from light, followed by magnetic microbeads incubation for 10 minutes in the same condition. The live cells and magnetic microbeads mix was added to the column placed in the magnetic multistand for further separation. CD71/ CD309/E-cadherin cells were harvested after flushing out form the column. A magnetic bead removal treatment was performed before transferring the cells to the culture medium.

### Single-cell dissociation and single-cell library construction

Briefly, a single-cell dissociation was prepared following the protocol described earlier. 10,000 cells of each tissue were loaded for library construction. Single cell libraries were prepared using Chromium Next GEM Single Cell 3’ Reagent Kit v3.1 (10X Genomics) according to the protocol detailed by the manufacturer with use of 10X Controller (10X Genomics, CA). The premade libraries were sent out to Novogene (Sacramento, CA) for Next Generation Sequencing.

### Single-cell RNA-sequencing analysis

We adopted the Optimus pipeline 5.1.1 of the Human Cell Atlas (PMID: 29206104) for the alignment and quantification of the short reads. Briefly, the Optimus pipeline partitions the short reads into different samples based on the barcodes, then uses STARsolo alignment software (version 2.7.9a, PMID: 23104886) for the trimming of raw input, correction of the cell barcodes and Unique Molecular Identifiers (UMIs), and mapping and quantification of the short reads. For the mapping and quantification, we used mouse reference genome version GRCm38 and a mouse gene model from GENCODE version M23. We used the output matrices for the downstream analyses. We used DropletUtils 1.14.2 to distinguish the empty drops from the real cells (PMID: 30902100). We used “lower = 1000” parameter to specify the lower bound on the UMI counts. We used ScDblFinder 1.8.0 for the detection and removal of artificial doublets in the single-cell RNA-Sequencing (doi: 10.12688/f1000research.73600.1). Then we used Seurat 4.1.0 (PMID: 34062119) to remove cells that have either less than 200 or more than 3000 genes detected. Of note, we did not remove the cells based on their mitochondrial contents for the better tracing of dying cells.

Before the quantification of gene expression and identification of cell type specific marker genes, we used SCTransform 0.3.3 (PMID: 31870423) for the normalization of UMI counts, while regressing out the mitochondrial contents. Then we used Seurat for the initial identification of cell clusters, dimension reduction, and the projections using Uniform Manifold Approximation and Projection (UMAP). We used Seurat’s “FindAllMarkers” function to generate a list of cluster-specific marker genes. In doing this, we set the minimum fraction of the cells as 0.01, and the minimum fold difference (log scale) as 0.25. From the initial clustering, we could identify erythrocytes from different differentiation stages in both the lung and kidney datasets. These cells displayed unique gene expression profiles with high hemoglobin gene expression and low transcript complexity and formed distinct cell clusters. Thus, we filtered out those cells and repeated the above procedure again. Sub-clustering of the lung epithelial cells and kidney endothelial cells followed the same procedure.

For the cell type annotations of each cell population, we used the mouse single-cell marker gene lists from the CellMarker database (PMID: 30289549) as well as the kidney-specific cell-type markers described in a single-cell RNA-Seq analysis of the mouse kidney (PMID: 31118232). We performed Fisher’s exact test between those known markers and the markers that we identified for each cluster, then adjusted the p values using the Bonferroni correction. We report the cell-type assignment that has adjusted p values less than 0.05 in this study, except for the kidney clusters K2, K5, K13, and K14, as well as the lung cluster L9, which we manually assigned cell types based on the differential expression analysis results.

### Lipidomics analysis

After pulling down the cells using magnetic microbeads (10^6 ∼10^7 cells), the cells were subjected to ultrasonic wave disruption in an ice bath at 50W for 3 minutes. 1.5mL Chloroform: MeOH (2:1, v/v), 0.5 mL ultrapure water was added to the cells and vortexed for 1 minute, followed by centrifugation for 10 minutes at 3,000 rpm at 4°C. Then we transferred the lower phase to a new tube and shipped the samples out in a dry ice bath. Mass spectrometry-based lipidomics and bioinformatics analysis were conducted by Creative Proteomics (Shirley, NY).

Lipidomics of oxidized lipids in the pulldown cells were performed in the Metabolomics Center at University of Aberta, Canada. All chemicals and reagents were purchased from Sigma-Aldrich (Markham, ON, Canada), except those specifically stated. LC-MS grade water, acetonitrile, methanol and formic acid were purchased from Fisher Scientific (Ottawa, ON, Canada). Lipids internal standard mixture, LipidRep, for sample normalization were purchased from Nova Medical Testing Inc. (Edmonton, AB, Canada).

The LC-MS analyses were performed by strictly following the SOP in both positive and negative ionization^44,45^. Sample extracts were injected between injection replicates of the QC pooled mixture prepared with the same sample batch. Parameters used for data acquisition are described below and summarized in the table. All LC-MS analysis were carried out on using Thermo Vanquish UHPLC linked to Orbitrap Exploris 240 Mass Spectrometer. The column used was Waters ACQUITY Premier CSH C18 column (100 × 1.0 mm, 1.7 µm particle size) and the column oven temperature was 50 °C. Mobile phase A was 10 mM NH_4_COOH in 40:40:20 MeOH/ACN/Water. Mobile phase B was 10 mM NH_4_COOH in 85:10:5 IPA/ACN/Water. The gradient setting was: t = 0 min, 25% B; t = 10 min, 53% B; t = 18.0 min, 99% B; t = 19.1 min, 99% B. The flow rate was 60 and 40 µL/min. The injection volume was 1.0 µL for positive ionization and 1.5 µL for negative ionization.

For mass spectrometer, data-dependent acquisition MS/MS scan mode was used to acquire both MS and MS/MS data for all samples. The isolation window was 1.2 m/z with resolution of 45000 for MS/MS acquisition, and resolution of 90000 for MS acquisition, with an m/z range from 150 to 1500. For both positive and negative ionization, MS/MS collision energies ramping from 15 eV to 30 eV were used. The detailed parameters for MS ESI source tuning can be found on the table.

### Quantitative real-time Polymerase Chain Reaction

Total RNA was extracted from cells using TRIzol reagent. After RNA purity and concentration measurement, reverse transcription was performed using iScript Reverse Transcription Kit according to the manufacturer’s instruction. Quantitative Real-time Polymerase Chain Reaction was carried out in a StepOne Plus PCR system using SYBR Green. The mRNA level of targeted genes was normalized to β-actin as an internal control. A detail about the qPCR primers is presented in supplementary information.

### Mouse lung stereology

The stereology is performed as described^46,47^. Mice lung from postnatal day 3 were harvested following isofluorane administration. The lungs were dissected under a dissecting microscope. The thoracic cavity was exposed by midsternal thoracotomy and dissection of the diaphragm at the point of contact with the ribcage. Tracheostomy was undertaken with a 22G blunt end cannula. The lungs were inflated with intratracheal instillation of air at an airway pressure of 20 cm H_2_O. Lungs were considered fully inflated once no visible increase in lung expansion was noted. The tracheostomy needle was partially withdrawn and the trachea ligated with a suture. The lung was gently removed and added to 4% PFA in PBS and incubated overnight at 4C with shaking. Following fixation, the lungs were embedded in 2 % agar and agar blocks sectioned at 3 mm thickness. The sections were stained using Richardson’s staining and imaged. Three separate litters were harvested and genotyped for GPX4 Tg and Cre+GPX4 mice. Quantification of airway spaces from a minimum of 3 mice pups per genotype using Stereo Investigator software, version 2022.2.3 (MBF Bioscience). The lung alveolar area and mean linear intercept was obtained by counting multiple fields in each section and data obtained after averaging each field of measurement from each genotype of mice.

#### Location of hGPX4 insert in mouse genomic DNA of GPX4 Tg mouse model

To identify the genomic location of hGPX4 insert in GPX4 Tg mice, we performed Illumina-based short-read DNA sequencing and Oxford Nanopore Technology long-read Direct sequencing. (1) Illumina-based short-read sequencing. We mapped the paired-end DNA-Seq reads to a custom reference containing (a) the whole mouse genome sequence (Genome Reference Consortium Mouse Build 39, GRCm39), (b) the human GPX4 gene sequence with its 100 bp upstream and downstream (chr19: 1,003,982-1,206,791 in the human genome reference GRCh38), and (c) the P1 vector sequence (GenBank ID: U09128). We used the Burrows-Wheeler Alignment tool (BWA) version 0.7.17 for mapping (PMID: 19451168). We used Delly version 1.1.8 (PMID: 22962449) for structural variant detection with parameters -q 20 (for mapping quality) and -s 1000 (minimum size of deletion call). In doing so, we filtered out interchromosomal sequencing pairs covering both the P1 vector and the region surrounding mouse chr1:78,585,525. This is because the latter contains a long poly-G sequence, which is also a known artifact of the sequencing platform (Nova-Seq, Illumina) that we used. (2) Oxford Nanopore long-read DNA sequencing. We used Dorado version 0.8.0 (https://github.com/nanoporetech/dorado) with its High Accuracy Model (HAC) for base calling, trimming, and mapping of the long reads. We used the same reference sequence (mouse genome + human GPX4 gene region + P1 vector) as for the short-read sequencing. We used Delly to detect if there was an insertion of hGPX4 or P1 vector into the mouse genome with the parameters “lr” and “-y ont”. Finally, we used Flye version 2.9.5 (PMID: 30936562) for de novo assembly of the human GPX4 or P1 vector-containing DNA fragments. We used “--nano-hq” for assembly and “--iteration 10” parameter to polish the output sequence, resulting in a 69,399 bp contig that is circular.

### Lipid Peroxidation (MDA) Assay

Lipid peroxidation assay was conducted following manufacturer’s instructions. Briefly, 1 mg amount of tissue was washed with ice cold PBS. We homogenized the tissue in lysis solution (Buffer + BTH) with a dounce homogenizer on ice. We then centrifuged the samples to remove the pellets and transferred the supernatant to a new tube. After loading both standard and samples to the microplate, we measured absorbance immediately at OD 532 nm for colorimetric assay and calculated the concentration of MDA according to the protocol provided by the manufacturer.

### Urine and serum analysis

To measure urine microalbumin levels, Albuwell M enzyme linked immunosorbent assay (ELISA) kit was used following the manufacturer’s instruction. Then the microalbumin value was normalized with urine creatinine, which was determined by the Creatinine companion kit with the same sample. The urine albumin excretion rate was calculated as the urine albumin/creatinine ratio (UACR). Serum urea nitrogen level was detected using Urea Nitrogen Colorimetric Detection Kit according to the manufacturer’s protocol.

### Image quantification and Statistical Analysis

Fluorescence was monitored in cells expressing specific signals and analyzed using Image J or Photoshop. In the lung H&E staining, we used Image-Pro-Plus 6.0 to assess dense/cystic burden in each section as well as the alveolar area and septal thickness. In the kidney H&E staining, we used Image-Pro-Plus 6.0 to assess hypertrophy/cystic glomerulus in the sections.

All quantitative data were presented as means± standard errors (SEs). Wilcoxson, Mann-Whitney U or Student’s *t*-test was applied to two-group comparisons. One-way or Two-way ANOVA was used for comparisons including more than two groups and *Tukey* test was used to estimate the significance between groups once significant differences were found. Differences were considered statistically significant when *P* < 0.05. The Chi square test and Fisher Exact test were used to estimate the significance of lung and kidney defects incidences. All statistical analysis were performed in GraphPad Prism software V8.

## Data Deposition and Code Availability

The single-cell RNA-Seq data is available on the GEO website (https://www.ncbi.nlm.nih.gov/geo/) under accession number **GSE210840**. An PDF file containing the values that were used to create all the graphs in the paper, is included in the paper in Data S1.

We deposited the metabolomics data to the EMBL-EBI MetaboLights^3^ database (DOI: 10.1093/nar/gkad1045, PMID:37971328) with the identifier MTBLS11771. Once the status of the submission is updated as ‘Public’, the complete dataset will be accessed on https://www.ebi.ac.uk/metabolights/MTBLS11771

## Conflict of interest statement

The authors have declared that no conflict of interest exists

### Subheading

Ferroptosis in embryonic development

## Supplementary figure titles and legends

**Figure S1.**
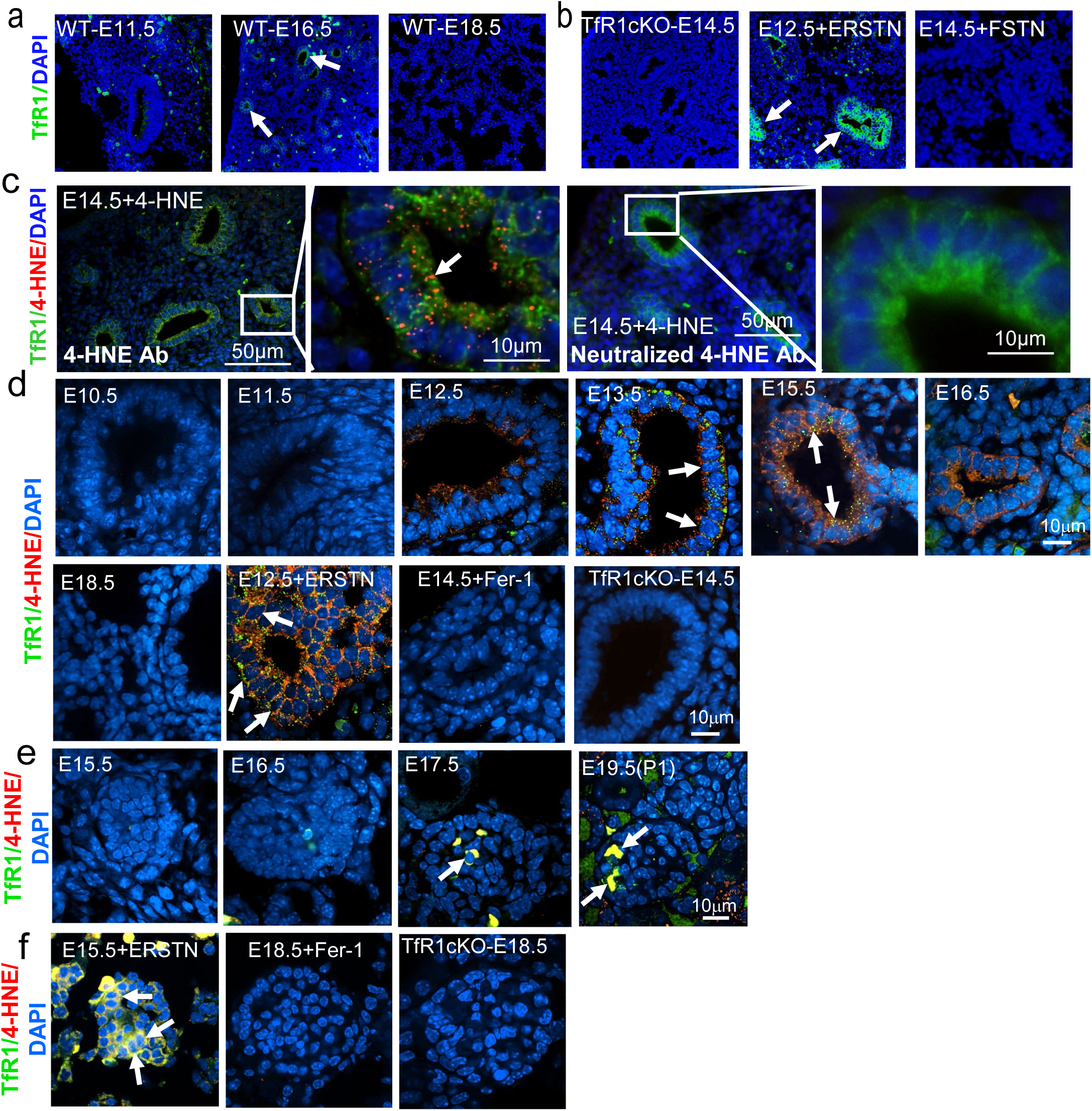
Presence of TfR1 single-positive and TfR1/4-HNE double-positive cells in embryonic lung distal epithelium and kidney glomeruli. (**a**) Representative immunostaining of TfR1 in development stages of lung from E11.5, E16.5 and E18.5.(**b**)TfR1 immunostaining for biological negative (TfR1 KO), experimental positive (E12.5 lung treated with the ferroptosis inducer Erastin (ERSTN)) and Negative (E14.5 lung treated with the ferroptosis inhibitor Ferrostatin-1 (Fer-1Fer-1)) control. (**c**) Representative immunostaining of double TfR1 and 4-HNE staining: negative control of 4-HNE using antibody neutralization by 4-HNE in E14.5 lung. **(d)** TfR1 and 4-HNE double immunostaining in different stages of lung development from E10.5 to E18.5 (upper panel) and biological negative (TfR1 KO), experimental positive (E12.5 lung treated with ferroptosis inducer ERSTN) and Negative (E1.5 lung treated with Fer-1) control (lower panel). **(e)** TfR1 and 4-HNE double immunostaining in different stages of embryonic kidney development from E15.5 to E19.5 (P1) (upper panel) and biological negative (TfR1 KO), experimental positive (E15.5 kidney treated with ERSTN) and Negative (E18.5 kidney treated with Fer-1) control (lower panel).

**Figure S2.**
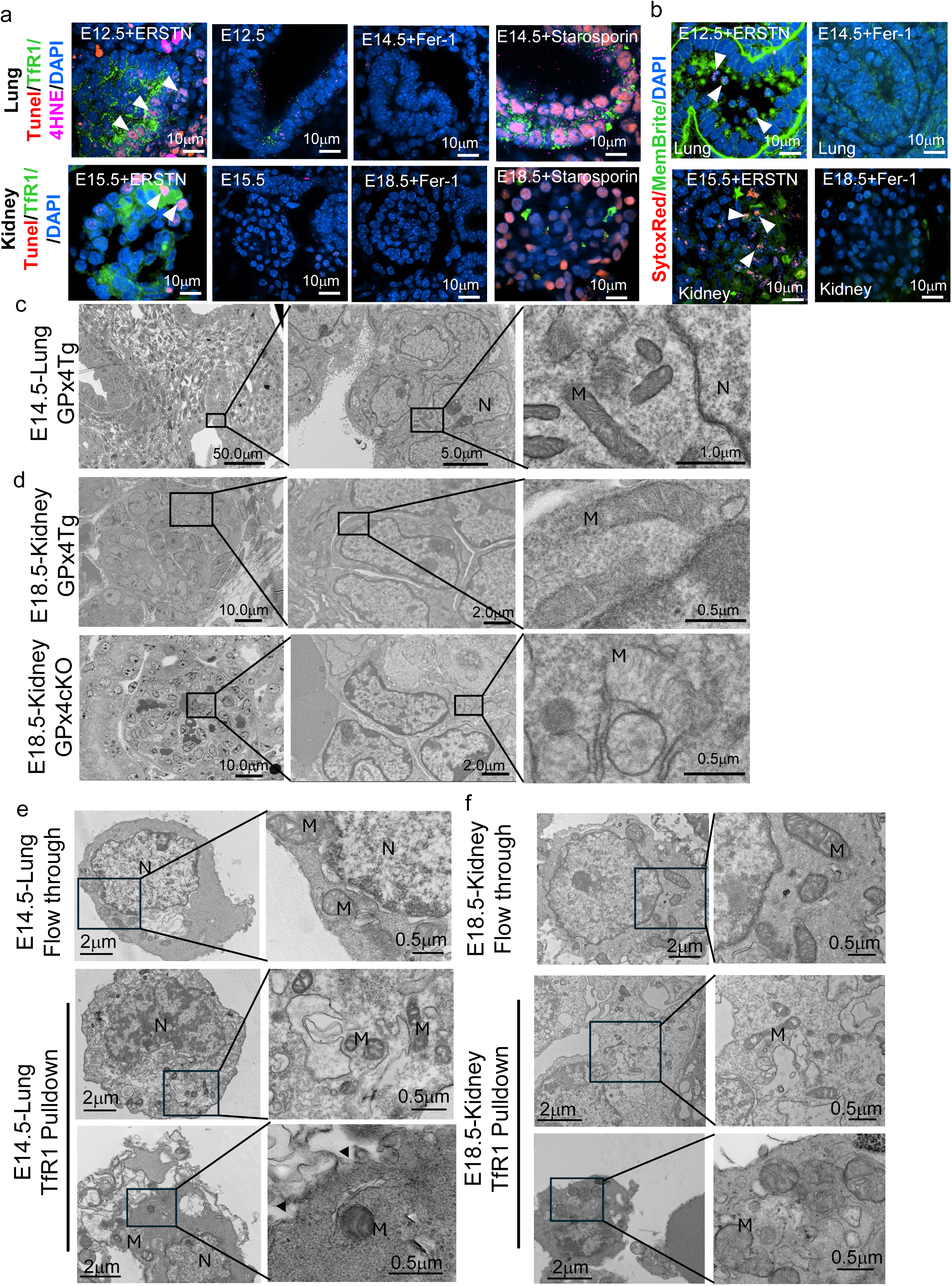
Presence of TfR1-4-HNE-TUNEL triple-positive cells, dead cells and mitochondrial defects in embryonic lung distal epithelium and kidney glomeruli cells. (**a**) Representative immunostaining of triple TfR1, 4-HNE and TUNEL staining in experimental positive and negative control for ferroptosis and TUNEL staining. The top panel stands for lungs and the lower panel for kidney. Experimental positive control for ferroptosis was the treatment of ERSTN (Erastin) and the negative control was the treatment of Fer-1 (Ferrostatin-1). TUNEL positive control was Staurosporine treatment. (**b**) Cell membrane dye MemBrite and nuclear dye SytoxRed perfusion staining of experimental positive (E12.5 lung treated with ERSTN) and Negative (E14.5 lung treated with Fer-1) control for lung (upper panel) and kidney (E15.5 kidney treated with ERSTN and E18.5 kidney treated with Fer-1) (lower panel). (**c**) Apreo SEM micrograph of E14.5 lung from GPX4 Tg and (**d**) E18.5 kidney glomerulus from GPX4 Tg (upper panel) and GPX4 cKO(GPx4ff:flk1 Cre) (lower panel). Mitochondria from specified region where ferroptotic cells are identified are focused. (**e**) Representative transmission electron microscopy images for flow-through and TfR1 pulldown cells from E14.5 lung and (**f**) E18.5 Kidney. Condensed nucleus (N), Shrink or swollen mitochondria (M) are indicated. Embryos from three different litters (*n* = 3) were analyzed.

**Figure S3.**
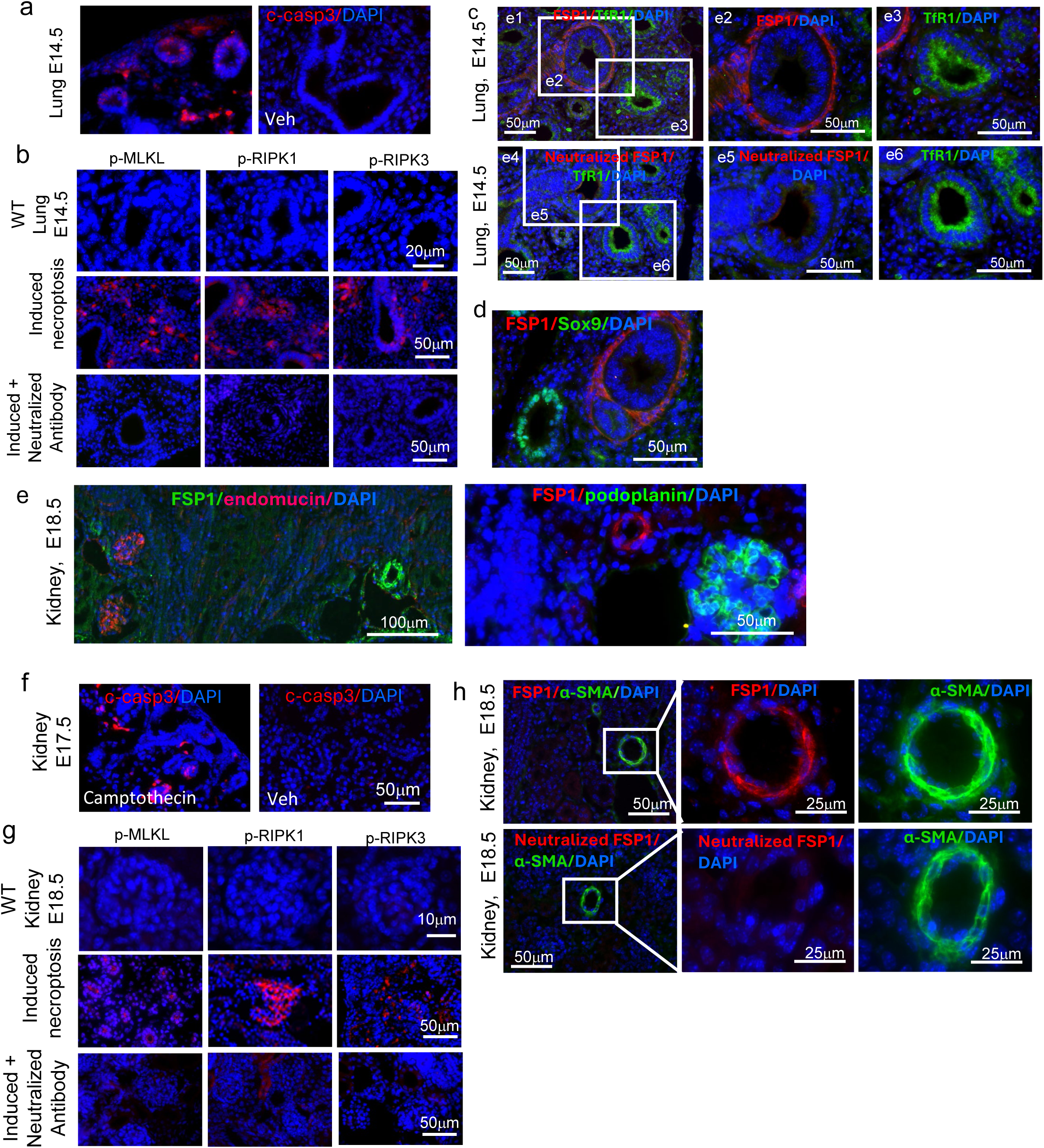
Verification of antibody specificity for immunofluorescence labeling and necroptosis and apoptosis do not occur in the lung and kidney during these stages of development. Lung tissues from E14.5 were treated with an apoptosis inducer, Camptothecin. A significant number of cleaved caspase-3 positive apoptotic cells were visible in the apoptosis-induced tissues but not in the controls (vehicle treated) (**a**). Necroptotic cells in the developing lung (**b**). Necroptotic cells were detectable only in the lung tissues treated with a necroptosis inducer. P-MLKL, p-RIPK1, and p-RIPK3 antibodies were neutralized with their corresponding proteins, then used in immunofluorescence staining, showing induced necroptosis was no longer detected by the neutralized antibodies. Dual labeling of E14.5 lung with FSP1and TfR1 antibody, showing no overlapping in labeling (**c**). After FSP1 antibody was neutralized with FSP1 protein, FSP labeling became negative (**c**). Dual labeling of FSP and Sox9 showed no co-localization in E14.5 lung (**d**). In the E18.5 kidney, labeling of FSP1 together with endomucin, podoplanin, or α-SMA indicated that FSP positive labeling co-localized with α-SMA positive vessels but not with endomucin or podoplanin positive glomeruli (e). A number of cleaved caspase3-positve cells were detected in E17.5 kidney treated with apoptosis inducer (**f**). There are no necroptotic cells observed in the developing kidney, and necroptosis staining was specific (**g**). Double staining of FSP1 and α-SMA in E18.5 kidneys (**h**).

**Figure S4.**
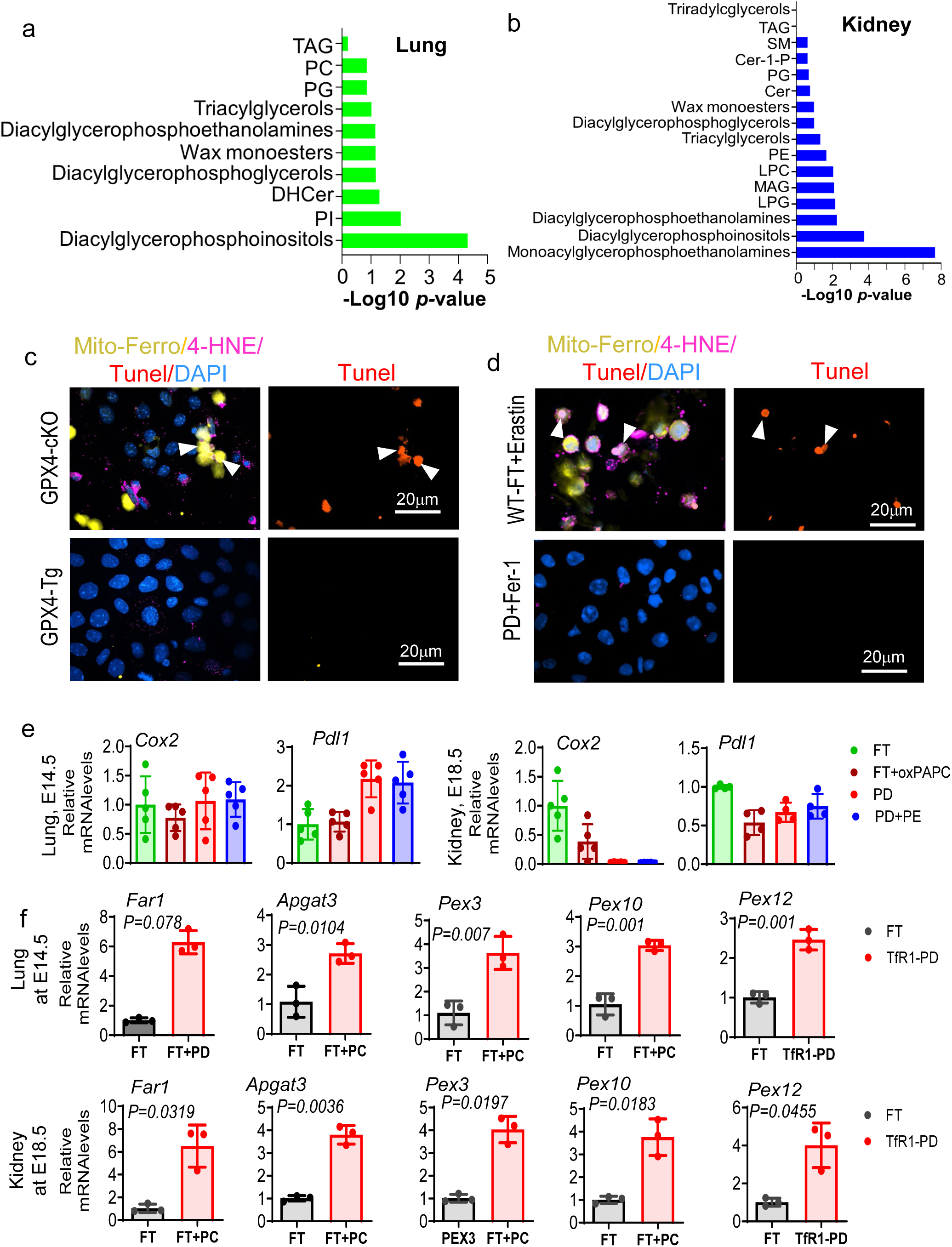
Lipidomics pathways enrichment, Mito-Ferro, 4-HNE and TUNEL positivity in TfR1 cells and peroxisomal biogenesis gene expression in the developing lung and kidney. (**a**) Pathway analysis for different detected lipids between the TfR1 pull-down group and the flow-through in E14.5 lung. (**b**) Pathway analysis for different detected lipids between the TfR1 pull-down group and the Flk1 pull-down group in E18.5 kidney. (**c**) Mito-FerroGreen, 4-HNE and TUNEL staining in cells from E14.5 GPX4 cKO (upper panel) and GPX4 Tg lungs (lower panel). (**d**) Experimental positive (Flow-through (FT) cells from E14.5 lung treated with ERSTN) and negative (TfR1 pulldown (PD) cells from E14.5 lung treated with Fer-1 (Ferrostatin-1)) for Mito-FerroGreen, 4-HNE and TUNEL staining are presented in upper panel and lower panel, respectively. (**e**) Real-time PCR analysis of genes including *Cox2* and *Pdl1* in flow-through (FT), TfR1 pulldown (PD), FT treated with oxPAPC and PD treated with the reduced lipid PE (PE (22:1_22:6)+Na) for the cells from E14.5 lung and E18.5 kidney. (**f**) Real-time PCR analysis of peroxisomal biogenesis genes including *Far1, Agpat3, Pex3, Pex10,* and *Pex12* in TfR1 pull-down and flow through cells using E14.5 lung (upper panel) and E18.5 kidney (lower panel). Embryo from 3 different litter (*n* = 3) (each liter includes 8-9 embryo) were used to prepare cells for TfR1 pull down. *P* values from Student’s *t*-test (**e, f**).

**Figure S5.**
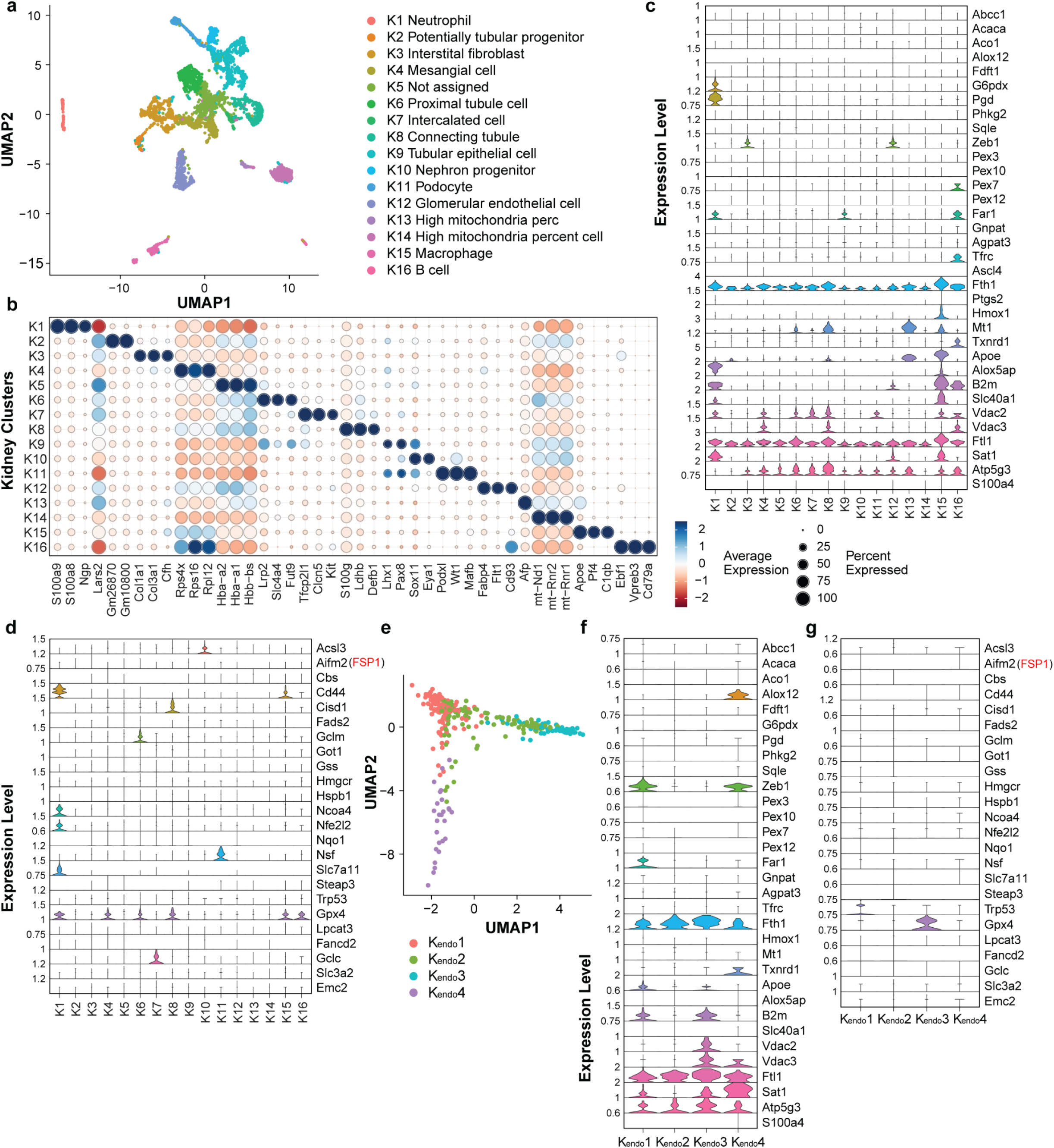
Single-cell RNA-sequencing confirms ferroptotic cells in the developing kidney. (**a**) Clustering plot of kidney cells (*n* = 6257) by UMAP (Uniform Manifold Approximation and Projection) based on the expression of highly variable genes. (**b**) Dot plot of cluster enriched gene expression. (**c**) Violin plots of ferroptosis positive gene arranged by clusters. (**d**) Violin plots of ferroptosis negative gene arranged by clusters. (**e**) Trajectory analysis of four sub-clusters cells from cluster 12. Violin plots of ferroptosis positive gene (**f**) and negative gene (**g**) arranged by sub-clusters.

**Figure S6.**
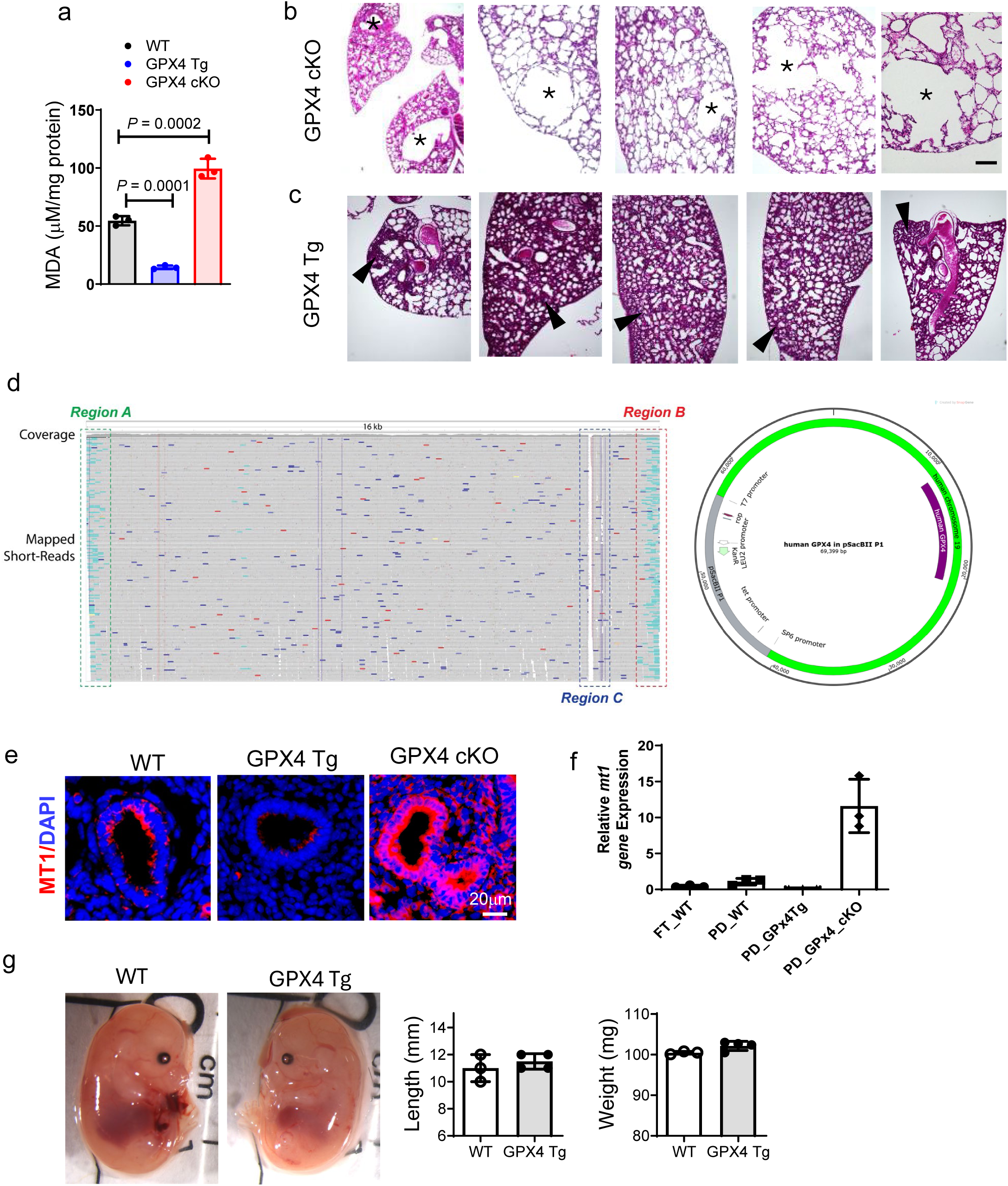
Lung defects in *Gpx4* conditional knockout and transgenic mice. **(a)** The lipid peroxidation (MDA) of E14.5 lungs was measured in WT, *Gpx4* TG and *Gpx4* cKO groups. **(b)** Representative H&E staining of *Gpx4* cKO lung at P3; asterisks indicate the cystic structure or disorganized alveoli. 12 lungs from 12 cKO embryos of 7 litters were analyzed. (**c**) Representative H&E staining of *Gpx4* TG lung at P3; black arrowheads indicate dense patches with minimal airspaces or thickening alveoli. 24 lungs from 24 Tg embryos of 7 litters were analyzed. (**c**) The lipid peroxidation (MDA) of E14.5 lung was measured in WT, *Gpx4* TG and *Gpx4* cKO groups. (**d**) Mouse genomic mapping for the *Gpx4* transgene in GPX4 Tg mice. (**e**) MT-1 immunostaining for E14.5 lung distal epithelia from WT, GPX4 Tg and GPX4 cKO mice. (**f**) Real-time quantitative PCR analysis of *Mt1* in E-Cadherin pulldown (PD) and flow-through (FT) cells using E14.5 lungs from WT, GPX4 Tg and GPX4 cKO. (g) *Gpx4* Tg does not affect embryo lengths nor weight. 3 embryos in WT and 4 embryos in *Gpx4* Tg group. ** indicates significant difference in one-way ANOVA followed by *Tukey* test (**a**).

**Figure S7.**
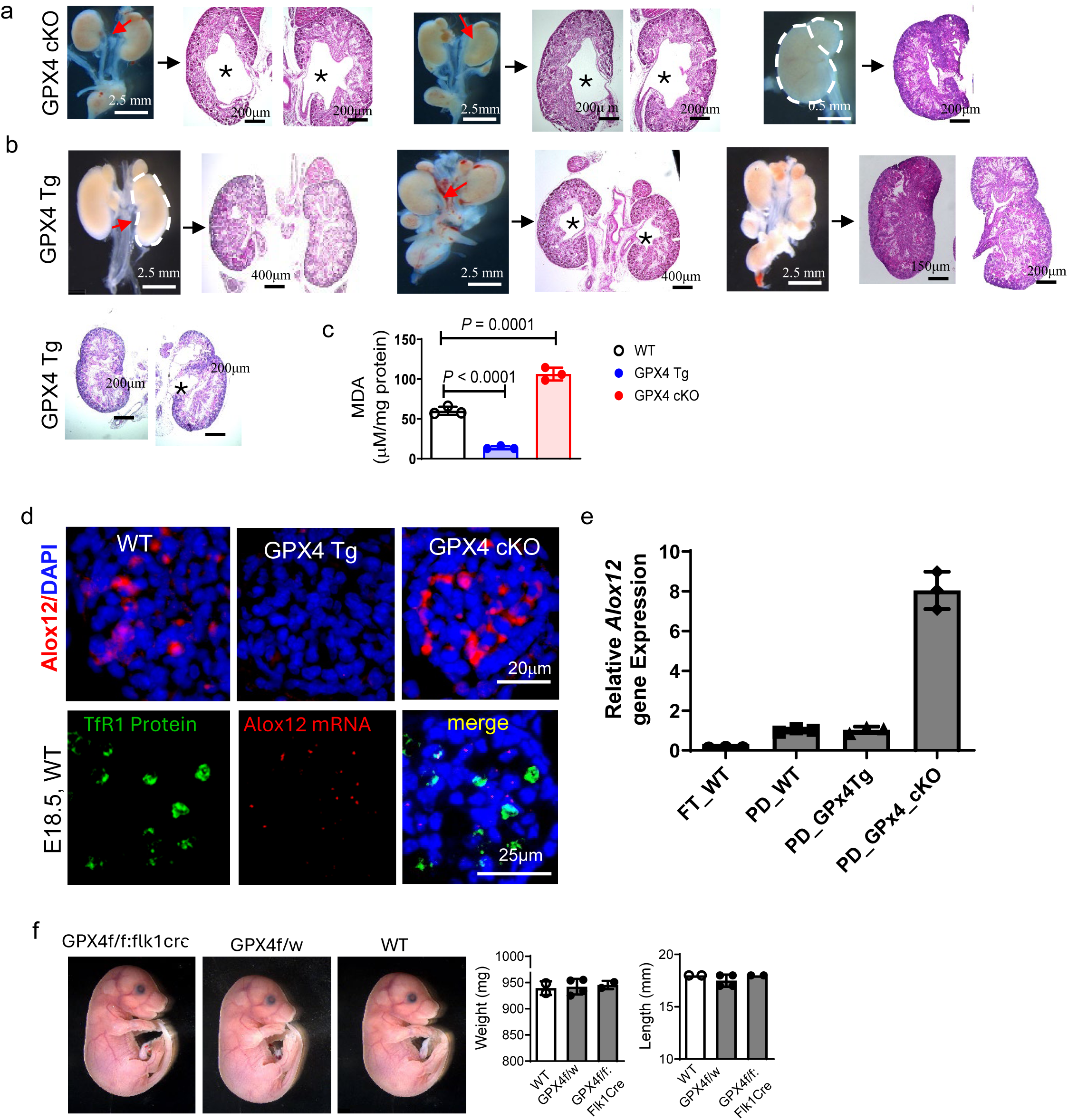
Kidney defects in *Gpx4* conditional knockout and transgenic mice. (**a**) Gloss morphology and representative H&E staining of *Gpx4* cKO kidneys; red arrows indicated the hydroureteronephrosis/hydronephrosis; dotted line lights up the duplex kidney; asterisks indicated the enlarge renal pelvis, indicating the hydroureteronephrosis or hydronephrosis in the kidney. 14 kidneys from 7 cKO embryos of 5 litters were analyzed. (**b**) Gloss morphology and representative H&E staining of *Gpx4* TG kidneys; red arrows indicated the hydroureteronephrosis/hydronephrosis; dotted line lights up the duplex kidney; asterisks indicated the enlarging renal pelvis, indicating the hydroureteronephrosis or hydronephrosis in the kidney. 52 kidneys from 26 Tg embryos of 7 litters were analyzed. **(c)** The lipid peroxidation (MDA) of E18.5 kidney was measured in WT, *Gpx4* TG and *Gpx4* cKO groups. **(d)** RNAscope dual staining (*Alox12* RNA Red; TfR1 protein green) (upper panel), and Alox12 immunostaining in in E18.5 kidney glomeruli cells from WT, GPX4 Tg and GPX4 cKO mice. **(e)** Real-time quantitative PCR analysis of *Alox12* in Flk-1 pulldown (PD) and flow-through (FT) cells of E18.5 kidneys (*n* = 3). (**f**) *Gpx4* cKO does not affect the embryo length and weight at E18.5. 2 embryos in WT and *Gpx4* cKO group and 4 embryos in *Gpx4*^f/w^ group. *P*<0.05 indicates significant difference in one-way ANOVA followed by *Tukey* test (**c**).

